# Engineered bacteria titrate hydrogen sulfide and induce concentration-dependent effects on host in a gut microphysiological system

**DOI:** 10.1101/2023.05.16.538950

**Authors:** Justin A. Hayes, Anna W. Lunger, Aayushi S. Sharma, Matthew T. Fernez, Abigail N. Koppes, Ryan Koppes, Benjamin M. Woolston

## Abstract

Hydrogen sulfide (H_2_S) is a gaseous microbial metabolite whose role in gut diseases is debated, largely due to the difficulty in controlling its concentration and the use of non-representative model systems in previous work. Here, we engineered *E. coli* to titrate H_2_S controllably across the physiological range in a gut microphysiological system (chip) supportive of the co-culture of microbes and host cells. The chip was designed to maintain H_2_S gas tension and enable visualization of co-culture in real-time with confocal microscopy. Engineered strains colonized the chip and were metabolically active for two days, during which they produced H_2_S across a sixteen-fold range and induced changes in host gene expression and metabolism in an H_2_S concentration-dependent manner. These results validate a novel platform for studying the mechanisms underlying microbe-host interactions, by enabling experiments that are infeasible with current animal and *in vitro* models.

## Introduction

Extensive research has linked dysbiosis in the gut microbiota with human disease^1–5^. In particular, microbiome composition is implicated in intestinal diseases such as irritable bowel syndrome (IBS), colorectal cancer, and Crohn’s disease^5–7^. Unfortunately, our mechanistic understanding of how specific microbes contribute to disease lags behind^8^. One of the methods by which bacteria exert influence on host health is through their production and breakdown of small molecule metabolites that act as chemical messengers between microbes and host cells^3,9,10^. Hydrogen sulfide (H_2_S), a metabolite produced by both host cells and the microbiota, is implicated in various gut diseases, but the underlying mechanisms are not understood and its role is debated^11–13^. Colonic H_2_S is produced by sulfate-reducing bacteria (SRB) and cysteine-degrading bacteria, with the latter being more abundant in the microbiome^14^. Several lines of evidence support a role for H_2_S as a toxic mediator of inflammation. Patients with inflammatory bowel disease (IBD) and ulcerative colitis (UC) present with higher SRB composition^15^ and fecal H_2_S levels^16,17^, respectively. *In vitro* studies have shown H_2_S donors induce cell cycle arrest in immortalized intestinal epithelial Caco-2 cells^18,19^ and affect proliferation of the immortalized intestinal epithelial cell line, HCT116, in a concentration-dependent manner^20^. Additionally, at high concentration, H_2_S inhibits colonocyte oxygen consumption and butyrate oxidation^11^, and intra-colonic injection of the sulfide donor NaHS leads to an up-regulation of inflammation-related genes^21^. Finally, colonization of genetically predisposed (IL10^−/−^) mice with the H_2_S producer *Atopobium parvulum* leads to a worsening of colitis, with the effect being reversed by treatment with bismuth, an H_2_S scavenger^22^. By contrast, other evidence points to H_2_S as an agent of stability at the mucosal interface^13^. In rats, H_2_S donors enhance gastric ulcer healing and colitis resolution^23,24^. In addition, inhibition of host H_2_S synthesis leads to increased susceptibility to mucosal injury, and low concentrations of H_2_S increase metabolic activity in colonocytes^20^. H_2_S’s role as a therapeutic or toxic mediator in each process is therefore contradictory, and appears to depend on concentration, the model system used, and the route of administration^11^. Further complicating the interpretation of these studies are the technical challenges associated with controlling and measuring this reactive, volatile metabolite in the intestine. Thus, a means to reliably control H_2_S concentration in the gut microenvironment would help to elucidate its concentration-dependent effects in human tissues.

To study microbially derived metabolites and their effects on the host, researchers have generally used two approaches: knocking out or overexpressing biosynthetic gene clusters in relevant bacteria and evaluating their impact *in vivo*; or introducing the metabolite in a static *in vitro* system (e.g. Transwell) by direct addition or the use of spent microbial media^8,25,26^. Both approaches present challenges. While animal models can recreate the complex biology of the gut, unfortunately findings often do not translate to human health with high accuracy due to species differences in anatomy and genetics. Additionally, their use poses ethical issues^27^. Mice colonized with a human microbiota are an advancement, but may not capture specific interactions in the human gut that have arisen from the co-evolution of the host and microbiota^28^. For static microbial/human cell co-cultures, the lack of fluid flow leads to microbial overgrowth in co-cultures, so only short time scales can be explored^29,30^. By contrast, *in vitro* flow systems, such as the gut-on-a-chip (GoC), are emerging as excellent models for studying host-microbe interactions. They are easier to probe than animal models, and recreate certain organ level functions like transport, digestion, and metabolism better than static Transwell and intestinal organoid cultures^31–33^. Further, GoC systems are under physiological flow, which is necessary for establishing complex microbiomes and enabling the study of host-microbe interactions^34–36^. Researchers have also begun to use GoC systems to test the activity of therapeutic microbes as an orthogonal preclinical model^37,38^. Recently, Nelson et al. recreated *in vivo* function of a synthetic probiotic for phenylketonuria with high accuracy in a GoC model, highlighting the utility of chips in bridging the *in vitro* to *in vivo* gap^38^. Further, chips enable direct manipulation and measurement of the gut metabolome, which led to the discovery of metabolites that activate the pathogenesis of enterohemorrhagic *Escherichia coli*^39^. This study would have been challenging in mice due to the difficulty in modulating and measuring individual metabolites along the digestive tract.

Previous GoC models have primarily been comprised of polydimethylsiloxane (PDMS), permeable to gas molecules^34–36,38,39^. Investigating the impact of gaseous mediators like nitric oxide (NO), carbon monoxide (CO), and H_2_S on gut homeostasis is not feasible using PDMS without surface modifications to reduce permeability, or using atmosphere-controlled incubators^40,41^. We have recently developed a cut & assembled method for manufacturing reconfigurable microphysiological systems (MPS), including gut-MPS (‘GMPS’ or ‘chip’), models out of thermoplastics^41,42^ to retain gas tension on the chip precisely for investigating gasotransmitters. We therefore hypothesized that our gas tension-retaining GMPS would provide the ideal opportunity to address hypotheses about H_2_S-host interactions that are infeasible with traditional model systems.

In this work, we engineered a library of microbial strains that reliably titer H_2_S across the gut physiological range, and used them to investigate the concentration-dependent effects of H_2_S on the host epithelium in our GMPS. Tunable production was achieved via titratable expression of L-cysteine (cysteine) desulfidase, which catalyzes the conversion of cysteine to H_2_S. Through a combination of gene knockouts, screening of heterologous and native desulfidases, different strength promoters, and transporter engineering, we generated a library of strains capable of titrating H_2_S across the gut physiological range (300-3,400 μM)^43^ in Hungate tubes. Our engineered strains were inoculated and stably colonized the GMPS for at least two days, actively metabolizing glucose and titrating H_2_S (75 – 1,200 μM) over the duration of the chip experiment. RT-qPCR analysis of epithelial cells showed an H_2_S concentration-dependent effect on host expression of *GADD45a*, a gene involved in DNA damage and cell cycle arrest known to be upregulated in the presence of H_2_S. Additionally, metabolic analysis of co-culture effluent from the lumen channel showed host oxidation of microbial H_2_S into thiosulfate. Finally, our platform allows for live visualization of colonization with confocal fluorescence microscopy. Overall, we have validated a powerful platform combining synthetic biology and GMPSs for investigating the role of H_2_S in the gut, which will support fundamental discoveries in human health.

## Results

### Gut microphysiological system (GMPS) characterization

The GMPS used herein is a derivative of the work by Hosic et al.^41^. The chip is laser cut and assembled layer-by-layer resulting in a chip with polymethyl methacrylate (PMMA) apical and basal channels separated by a polyester (PET) membrane (**Fig. 1a**). Constant perfusion of culture medium was similar to physiological flow (flow rate 3 μL/minute, shear stress 0.076 dyne/cm^2^) to mimic shear forces in the intestine^36,42^. After ten days, a homogeneous immortalized Caco-2 epithelial monolayer populated the entire length of the PET membrane (**Fig. 1b**). Caco-2 tight junctions were validated by immunofluorescent staining for tight junction protein Zonula Occludens-1 (ZO-1) and the live nuclei stain, Hoechst (**Fig. 1c**). This GMPS is fabricated from gas impermeable PMMA which retains gas tension better than PDMS^41,42^, enabling the study of H_2_S on host epithelial cells (**Fig. 1e**). Effluent media can be used for on-chip metabolic analysis and quantifying monolayer permeability. The GMPS provides easy access to the cell culture channels enabling RNA extraction from host cells (**Fig 1e**).

**Figure 1.**
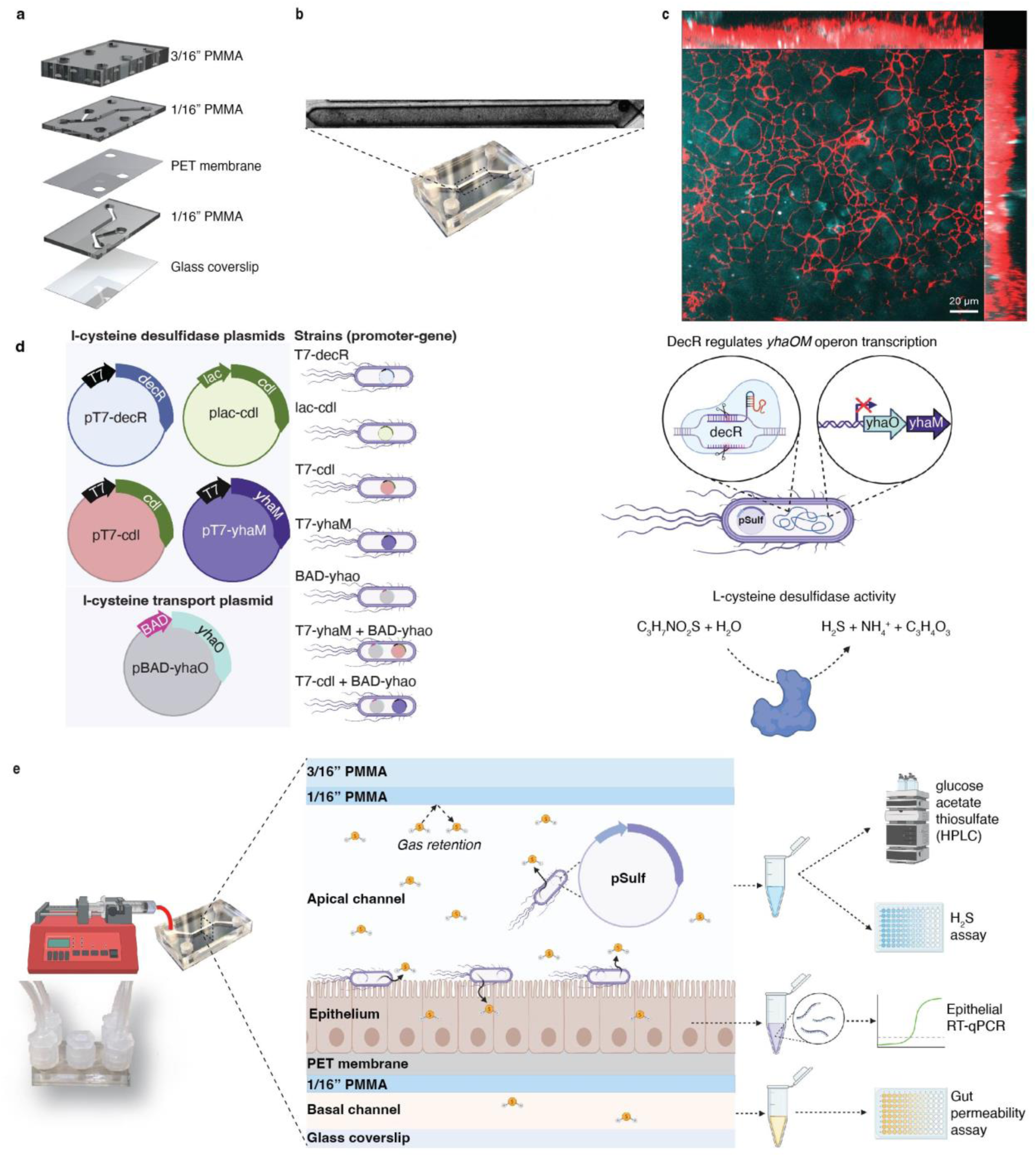
Characterization of GMPS and synthetic biology strategy. **a** Exploded view of the GMPS. Layers were adhered with double-sided 50 µm 3M adhesive tape. **b** Full channel brightfield image of Caco-2 cells in GMPS after ten days under flow. Multiple images were stitched together to create the tiled image. **c** Immunofluorescent staining of Caco-2 cells after ten days in GMPS. Immunostaining for ZO-1 (red) and Hoechst (cyan). The image is a z-stack maximum intensity projection. **d** Schematic of strain development and different promoter-gene combinations to control H_2_S titration. Different strength promoters lac, BAD, or T7, were used to control the expression of the desulfidases (*cdl* or *yhaM*) or the putative cysteine transporter, *yhaO*. DecR controls the *yhaOM* operon; *yhaO* is a putative cysteine transporter, and *yhaM* is a cysteine desulfidase. *decR* was deleted from MG1655 and used as the base strain moving forward. **e** Schematic of GMPS co-culture with engineered microbes and end-point analyses. Figure 1d,e were created with biorender.com.

### Strain development and cysteine desulfidase activity

Wild-type (WT) *E. coli* MG1655 possesses multiple cysteine desulfidase genes^44–48^ and produces 70 ± 41 μM and 1719 ± 97 μM in non-growth (no glucose or amino acids) and growth media, respectively (**Fig. 2a,b**). H_2_S levels in the intestine range from 300 to 3,400 μM^43^. To bring baseline H_2_S production to 300 μM, we individually deleted the *E. coli* desulfidases, *sseA* (also known as *mstA*) and *malY*, and the transcription factor *decR* (also known as *ybaO*)^44–50^. The *decR* gene codes for the transcription factor, DecR, which is activated by cysteine and regulates the *yhaOM* operon in *E. coli*^44^. *yhaO* codes for the putative cysteine transporter and *yhaM* a cysteine desulfidase (**Fig. 1d**). The effects on growth were assessed in M9 minimal media (**Supp. Fig. 1a**) and M9 minimal media with 5 mM cysteine (**Supp. Fig. 1b**) to test the mutants’ ability to metabolize cysteine. The *decR* mutant had slower growth kinetics, likely due to its reduced capacity to detoxify cysteine^44^. We then tested H_2_S production in non-growth and growth conditions (**Fig. 2a,b**). H_2_S production was an order of magnitude higher in growth conditions. Consistent with previous literature, the *decR* mutant reduced background H_2_S levels by approximately 85% (257 ± 50 μM, **Fig. 2b**). No reduction was seen for the *sseA* and *malY* strains. The literature is divided regarding the cysteine desulfidase activity of these gene products^44–46,50^, attributable perhaps to different *E. coli* strains, oxygen concentrations, growth media, or sulfide detection methods used in the various studies. Since it was critical the base strain produced below 300 μM H_2_S, the *decR* strain was selected as the chassis strain for subsequent engineering.

**Figure 2.**
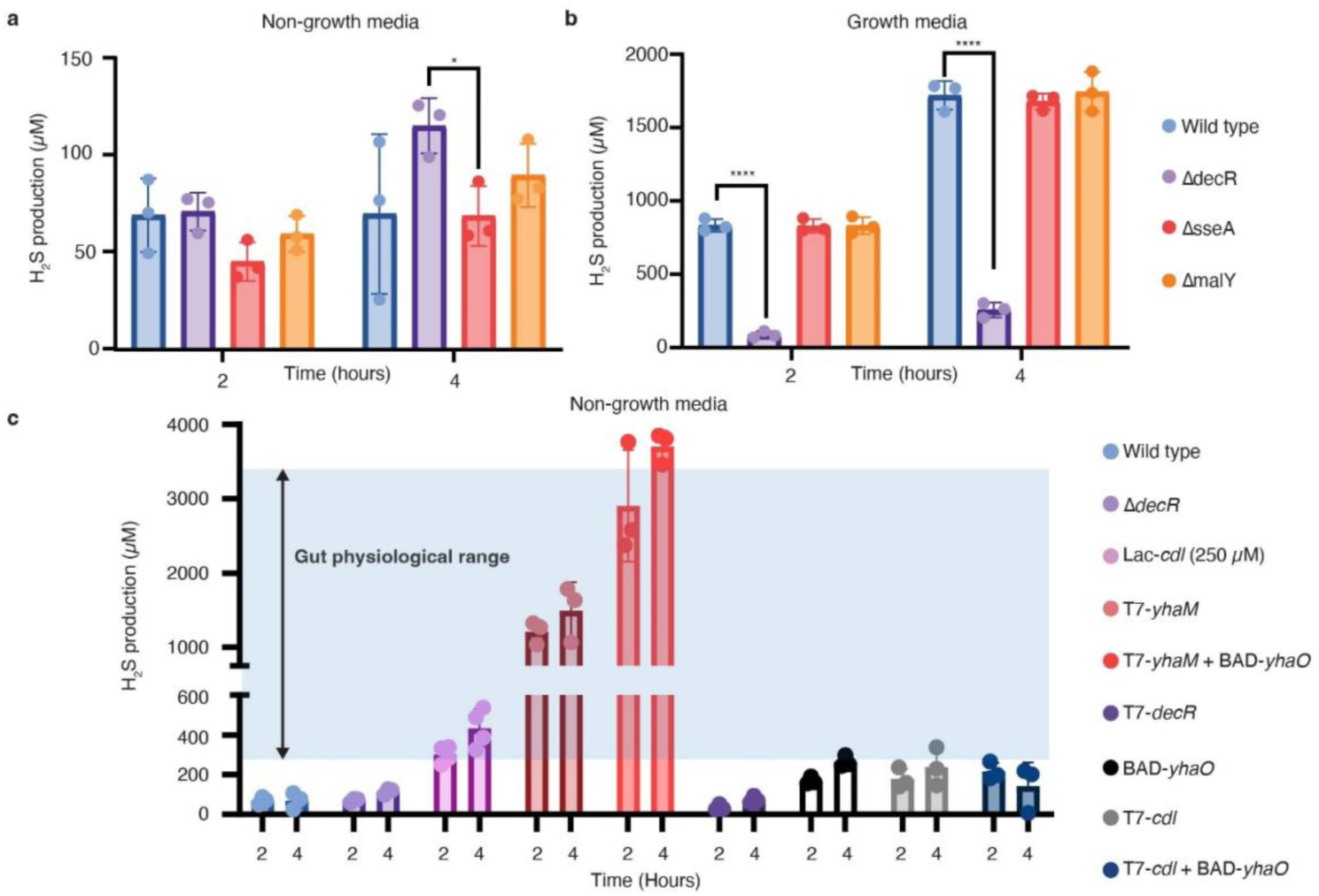
H_2_S production across the gut physiological range with engineered strains in Hungate tubes. **a-c** H_2_S production quantified in Hungate tubes over four hours, starting OD_600_ 0.4. Samples were taken while maintaining a closed system. **a** The three knockout strains were compared to wild type MG1655 in PBS++ (non-growth media) or **b** in M9 minimal media (growth media). **c** Engineered strains tested in PBS++ (non-growth media). The blue-shaded region represents the estimated gut physiological range (300-3,400 μM H_2_S). n = 3 independent experiments, for Lac-*cdl*; n = 4 independent experiments. Error bars represent SD, and bars represent the mean value. *p < 0.05, ****p < 0.0001. 2-way ANOVA with post hoc Tukey analysis using a 95% confidence interval. Source data can be found in the Source Data File.

To develop strains capable of controllably producing higher levels of H_2_S, we next assembled a small library of plasmids with various cysteine desulfidase homologs under the control of different promoters (**Fig. 1d**), then transformed them into the chassis strain. H_2_S production was tested in Hungate tubes – a closed system – to minimize H_2_S evaporation. When cysteine was omitted from the media, no H_2_S was produced (Source Data File). Strain Lac-*cdl* carrying the *Fusobacterium nucleatum* desulfidase, *cdl*, under control of the *lac* promoter produced increasing levels of H_2_S (200 ± 24 μM to 434 ± 96 μM) with increasing inducer concentration (**Supp. Fig. 2**). This was significantly below the upper bound of the physiological range, 3,400 μM. To improve H_2_S production, we explored overexpression of *decR* and the genes it regulates (the transporter *yhaO* and desulfidase *yhaM)* as well as further upregulation of *F. nucleatum cdl* with the phage T7 promoter (**Fig. 2c**). These strains produced H_2_S across the gut physiological range (blue shaded region in **Fig. 2c**).

Co-expression of *yhaO* and *yhaM* (T7-*yhaM* + BAD-*yhaO*) resulted in the highest levels of H_2_S, producing 3,707 ± 214 μM, 210% more H_2_S than the sum of the individual strains, T7-*yhaM* and BAD-*yhaO*. There was a negative effect when overexpressing *cdl* and *yhaO* (T7-*cdl* + BAD-*yhaO)*, reducing H_2_S production compared to the unique strains T7-*cdl* and BAD-*yhaO* (**Fig. 2c**). T7-*decR* in non-growth media did not show increased H_2_S production. We speculate this is because DecR is activated by cysteine^44^, which was added when cells were in the non-growth media, limiting transcription of *yhaO and yhaM*.

### Titratable microbial H_2_S production in the GMPS

Next, the engineered strains were inoculated in the GMPS to test their performance and production of H_2_S in a simulated gut environment (**Fig. 3a**). The GMPS has different physical conditions than Hungate tubes, including constant perfusion and shear stress which may affect microbial gene expression and activity^51^. Based on Hungate tube data (**Supp. Fig. 2**), the Lac-*cdl* strain (herein named ‘low-sulfide strain’) was used to demonstrate H_2_S titration by simply changing concentrations of the inducer, IPTG. Taking advantage of the high degree of experimental control over inlet streams perfusing the GMPS, 5 mM cysteine, varying amounts of inducer, and antibiotic(s) for plasmid maintenance were continuously fed for the duration of the experiment.

**Figure 3.**
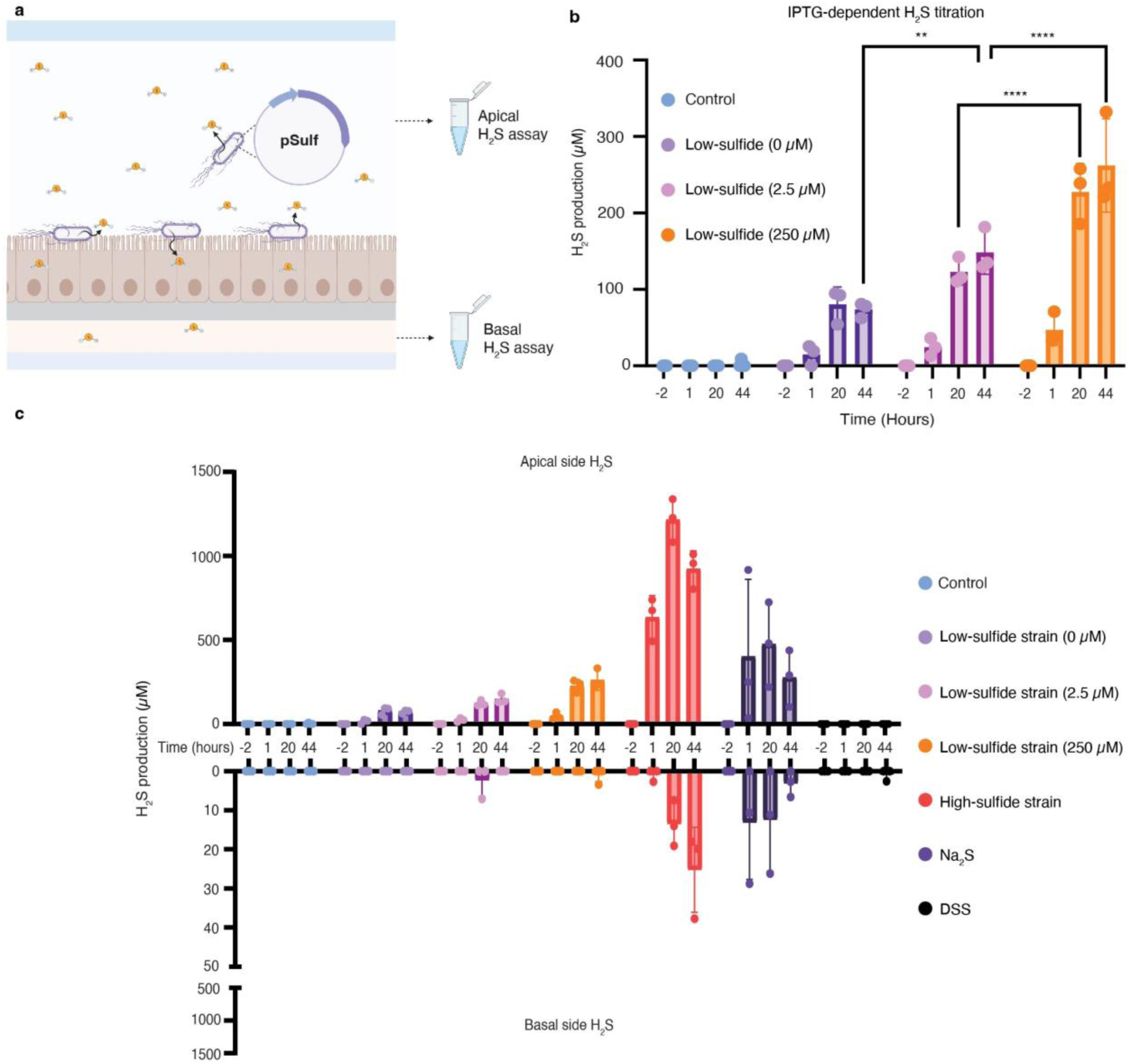
Titratable microbial H_2_S production in the GMPS. **a** Schematic of the experimental set-up. Cysteine was perfused through the chip, and H_2_S was produced by the engineered strains and collected for measurement with the methylene blue assay. **b-c** H_2_S was detected from the GMPS apical and basal effluents for two days. The effluent was collected in a tube containing zinc acetate, which effectively trapped H_2_S into stable ZnS. **b** Apical H_2_S production by the low-sulfide strain on chip by IPTG-dependent induction. Concentrations in parentheses correspond to 0, 2.5, and 250 µM IPTG induction. **c** Apical H_2_S production from all strains tested on the chip is shown in the top half of the graph. The bottom half of the graph shows H_2_S detected in the basal channel for each condition. Note the y-axes have different scales. The control chip has 5 mM cysteine, chloramphenicol, and 250 µM IPTG. n = 3 independent experiments, for Control; n = 4 independent experiments. Error bars represent SD, and bars represent the mean value. **p < 0.01, ****p < 0.0001. 2-way ANOVA with post hoc Tukey analysis using a 95% confidence interval. Source data can be found in the Source Data File. Figure 3a was created with biorender.com.

Samples were taken before strain inoculation to ensure there was no baseline H_2_S or microbial contamination in the GMPS (hour −2 in **Fig. 3**). After bacterial inoculation (hour 0), H_2_S was measured at hours 1, 20, and 44 (**Fig. 3a**). GMPSs containing no bacteria and only 5 mM cysteine, 250 μM IPTG, and chloramphenicol served as the negative control, confirming Caco-2 cells are unable to convert cysteine into significant levels of H_2_S. The low-sulfide strain produced across a 3.5-fold range (74 – 262 μM H_2_S) by varying inducer concentrations from 0, 2.5, and 250 μM IPTG (**Fig. 3b**). These strains permit a high degree of control over the lower range of H_2_S levels in the chip, enabling investigation of how small changes in H_2_S levels impact host biology. To access the upper bound of the gut physiological range, T7-*yhaM* + BAD-*yhaO* (herein named ‘high-sulfide strain’) was inoculated on the chip. In contrast to the precise H_2_S modulation of the low-sulfide strain, the high-sulfide strain maximized H_2_S production on the chip, producing 1,216 μM H_2_S (**Fig. 3c**). The lower production of H_2_S on chip compared to Hungate tubes could be explained by some evaporation during sample collection, or alteration of transcription or activity resulting from the constant perfusion and associated shear stress activity^51^.

We hypothesized localized sulfide production by microbes on-chip may provide different concentration profiles than the use of chemical sulfide donors, which are routinely used in animal models to investigate the effects of H_2_S in the intestinal environment^23,24^. To test this, we added 5 mM Na_2_S to the syringes perfusing the GMPS. Interestingly, effluent H_2_S levels had large standard deviations relative to the mean, with coefficients of variation of 1.143, 0.530, and 0.607 at hours 1, 20, and 44, respectively (**Fig. 3c**). Comparatively, the high-sulfide strain had lower coefficients of variation: 0.201, 0.105, and 0.116 at hours 1, 20, and 44, respectively. These results highlight the inconsistency and unpredictable nature of exogenously dosing a volatile metabolite, such as sulfide. Nearly 90% of the exogenously dosed H_2_S is unaccounted for and likely evaporated during experimental preparation and from the syringe, thus making it difficult to dose targeted concentrations. H_2_S can passively diffuse through cell membranes and is found in human serum. We therefore analyzed effluent from the basal channel, detecting H_2_S at levels similar to serum (22.2 – 81.0 μM)^52^ (**Fig. 3c**) in chips with the high-sulfide strain or exogenous Na_2_S. Together, these results suggest our system could be used to more accurately dose H_2_S through microbial production, and probe the systemic role of gut-derived H_2_S.

To assess microbial viability on-chip, apical and basal effluents were collected and used for colony counting. The engineered strains were viable on the GMPS for the full two days of the experiment (**Supp. Fig. 3**). The two-day co-culture did not result in translocation of the microbes. However, the low-sulfide strain was detected in the basal channel in one instance (n = 1), but at relatively low densities compared to the apical channel and only at the final time point (**Supp. Fig. 3**).

### Characterization of microbial colonization and metabolism in GMPS

Based on the steady H_2_S production and viable *E. coli* cells in the effluent over two days, it was plausible *E. coli* was colonizing the chip. The apical channel contained non-growth media and cysteine, and in Hungate tubes, cysteine alone was insufficient to support growth (Source Data File). We therefore hypothesized there may be nutrient diffusion from the basal to apical channel contributing to *E. coli* metabolism and colonization. Previous work in GoC systems demonstrated the enterorecirculation phenomenon^38^, a physiological process in which basal metabolites diffuse into the apical lumen, making it possible glucose could diffuse through the Caco-2 monolayer into the apical channel for *E. coli* utilization (**Fig. 4a**). To test this, we measured the levels of glucose and *E. coli* metabolites (acetate, formate, succinate) in apical effluent collected before and after inoculation of *E. coli*, and from a negative control chip with no microbe inoculation. At baseline, glucose was present in the apical channel at ~0.1 g/L. In negative control chips, no acetate was found, and glucose levels stayed constant for the duration of the experiment. After chips were inoculated with *E. coli*, glucose fell to undetectable levels, and acetate was produced at ~0.1 g/L. The GMPS was under flow which implies there was constant basal-to-apical glucose flux, and glucose was undetectable after microbial inoculation, indicating that *E. coli* actively metabolized glucose for two days in the co-culture (**Fig. 4b**).

**Figure 4.**
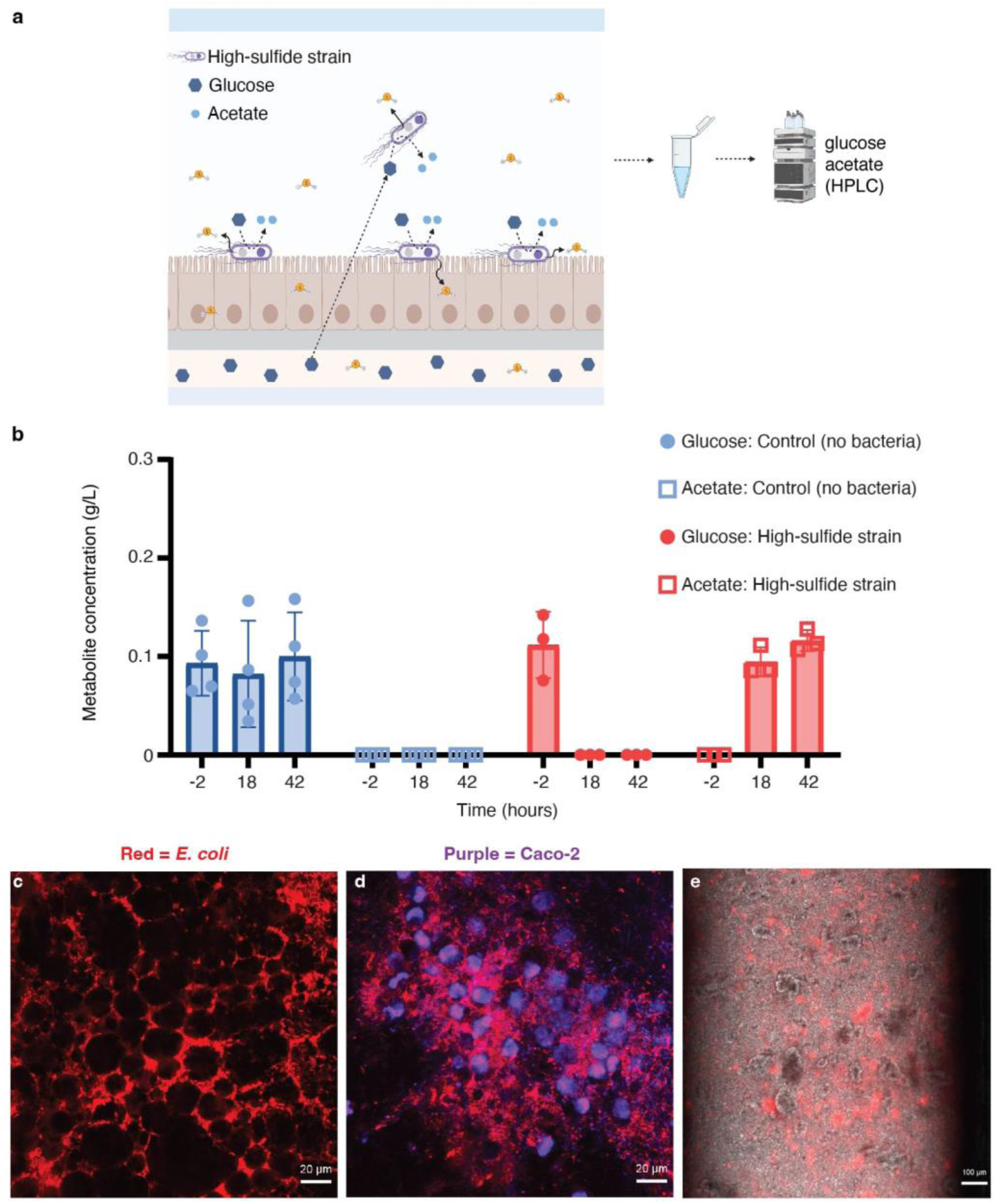
Characterization of microbial colonization and metabolism in GMPS. **a** Schematic of the experimental set-up. Glucose diffuses into the apical channel and is metabolized into acetate by *E. coli*. Cysteine was perfused through the chip, and the high-sulfide strain produced H_2_S. **b** Apical effluents collected from GMPS experiments were used for HPLC analysis. Metabolite levels were quantified before (hour −2) and after (hours 18 and 42) the treatment condition began, either negative control conditions or inoculation of the high-sulfide strain. Control; n = 4 independent experiments and high-sulfide strain; n = 3 independent experiments. **c-d** Representative images of the co-culture. 400X live confocal images, post-processed with Gaussian smoothing. **c** High-sulfide strain expressing *RFP* (red) in GMPS after two days of co-culture. **d** Z-stack cross-sectional image of the high-sulfide strain expressing *RFP* (red) and Caco-2 cell stained nuclei (purple) in the GMPS after one day of co-culture. **e** Representative image of the co-culture. 100X live image of RFP expressing *E. coli* MG1655 (red) with RFP and DIC filters captured with an inverted microscope. Error bars represent SD, and bars represent the mean value. Source data can be found in the Source Data File. Figure 4a was created with biorender.com.

To further analyze the extent of colonization and localization of cells within the GMPS, a chip inoculated with the high-sulfide strain engineered to constitutively express red fluorescent protein (*RFP*) was imaged by confocal microscopy (**Fig. 4c,d**). A video taken with an inverted microscope shows the GMPS was colonized by WT *E. coli* MG1655 expressing *RFP* under flow conditions for multiple hours (**Supp. Movie 1**) (**Fig. 4e**). Imaging confirms these strains maintain a population on a chip under constant flow, without washout, for the duration of the experiment (two days). Taken together, the metabolic, microscopy, and plating data show a mix of colonization and planktonic bacterial growth in the GMPS, consistent with the *in vivo* microbiota organization^53^.

### Microbially derived H_2_S affects Caco-2 cells in a concentration-dependent manner

H_2_S is hypothesized to play a role in IBD onset and impaired epithelial barrier function^54^. To investigate the concentration-dependent effects of H_2_S on gut permeability, Lucifer Yellow, a paracellular permeability marker, was added to the syringes feeding the GMPS (**Fig. 5a**). The effects of H_2_S were investigated using negative control conditions, dextran sulfate sodium (DSS) as a positive control which increases cell permeability^55^, exogenous Na_2_S, low-sulfide strain, and high-sulfide strain (**Fig. 5a,b**). These data were collected during the same experiment and correspond to H_2_S data in **Figure 3c**. The positive control, DSS, significantly disrupted the monolayer compared to the control, similar to previous work in GoC systems^55^. This two-day experiment showed no significant change in apparent permeability (P_app_) between the control and high-sulfide strain. There was a significant difference in P_app_ between Na_2_S and control at hour 18, but this may be due to higher baseline P_app_ values at hour −4 for Na_2_S, attributable to Caco-2 passage number differences^56^.

**Figure 5.**
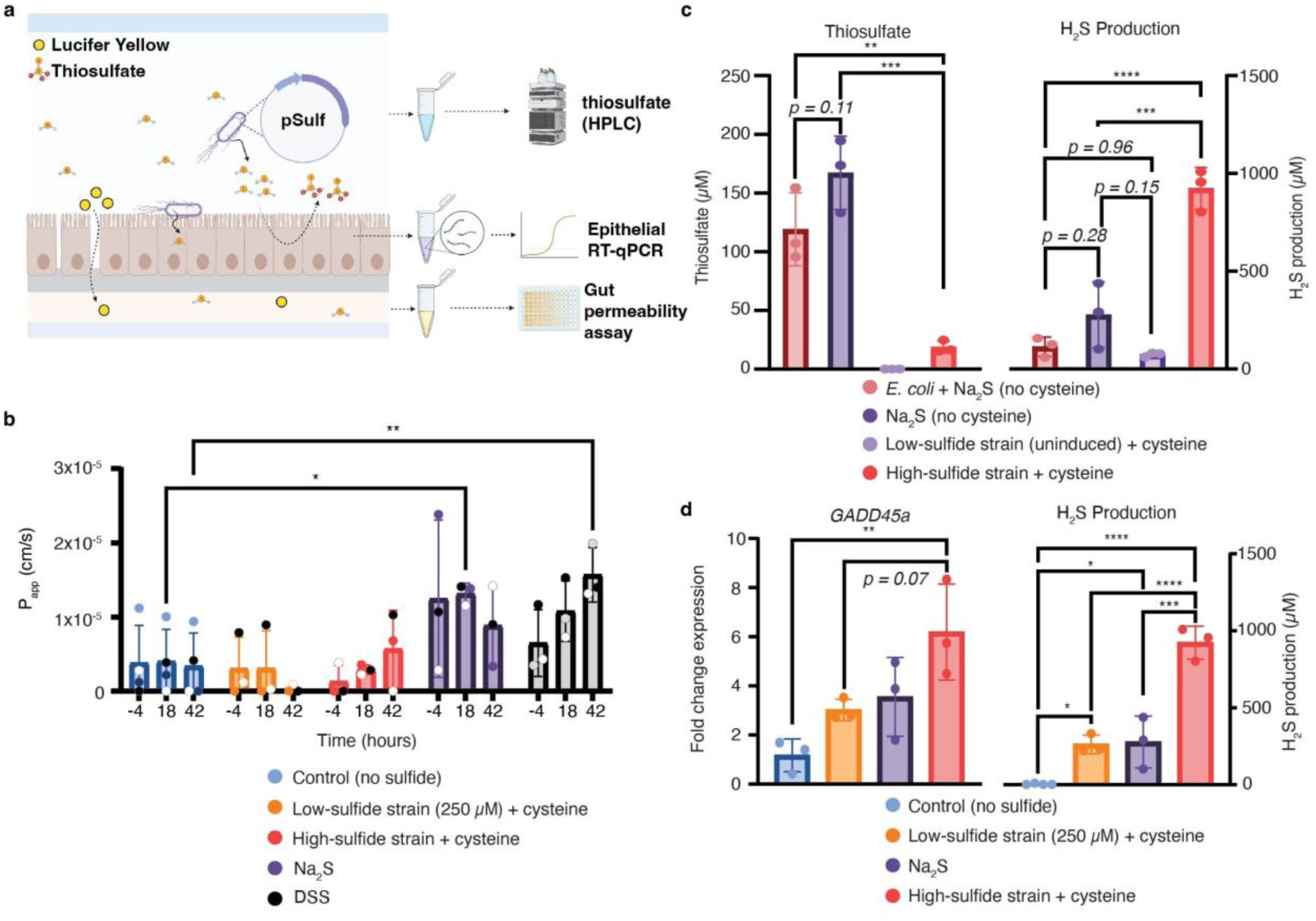
Microbially derived H_2_S affects Caco-2 cells in a concentration-dependent manner. **a** Schematic of the experimental set-up. Cysteine was perfused through the chip, and the engineered strains produced H_2_S. Paracellular diffusion into the basal channel was quantified using Lucifer Yellow, and P_app_ was determined. Microbially derived or exogenous H_2_S was metabolized into thiosulfate by Caco-2 cells and was detected with HPLC. **b** P_app_ (cm/s) was determined by collecting basal effluent before (hour −4) and during treatment conditions (hours 18 and 42). Colored dots indicate samples taken from the same biological replicate for identifying trends in permeability before and after treatment. n = 3 independent experiments, for Control; n = 4 independent experiments. **c** (left) Apical effluent was collected and analyzed for thiosulfate via HPLC, data shown from hour 42. (right) Corresponding apical H_2_S production levels at hour 44. *E. coli* + Na_2_S represents the high-sulfide strain (no cysteine) with exogenous Na_2_S. n = 3 independent experiments. **d** Caco-2 RNA was extracted after two days of treatment. (left) Fold change expression of *GADD45a* mRNA levels (reference gene *GAPDH*). Fold change expression was normalized to the Control. (right) Corresponding apical H_2_S production levels at hour 44. Error bars represent SD, and bars represent the mean value. *p<0.05, **p < 0.01, ***p < 0.001, and ****p < 0.0001. 1-way or 2-way ANOVA with a post hoc Tukey analysis using a 95% confidence interval. For panel b, all values were compared to the control group. Source data can be found in the Source Data File. Figure 5a was created with biorender.com.

We next examined crosstalk of sulfur metabolites by human and microbial cells. H_2_S can be oxidized by human sulfide quinone oxidoreductase (*SQOR*) and further metabolized into thiosulfate^57^. Thiosulfate in apical effluent was quantified with HPLC after derivatization with monobromobimane. As expected, thiosulfate was not detected in no-bacteria control (no sulfide) conditions, indicating cysteine was not converted to thiosulfate by Caco-2 (**Supp. Fig. 4**). Interestingly, with the high-sulfide strain, thiosulfate levels were much lower than on the Na_2_S chips (**Fig. 5c left**, *p* = 0.0002), despite the fact that the high-sulfide strain chips produced more H_2_S than Na_2_S-only chips (**Fig. 5c right,** *p* = 0.0003). In these experiments, Na_2_S was dosed on chips without colonized *E. coli*, thus one explanation for this contradiction is that the metabolically active bacterium might metabolize thiosulfate. Indeed, *E. coli* is known to be able convert thiosulfate through rhodaneses to sulfite and sulfide^58^. To explore this possibility, we measured thiosulfate production in a chip perfused with Na_2_S and inoculated with *E. coli*, but *without* addition of cysteine (i.e., no microbial H_2_S production). Thiosulfate levels were not significantly different between these conditions and the Na_2_S-only chip (**Fig. 5c left**, *p* = 0.11), suggesting *E. coli* is not a dominant consumer of thiosulfate in this system. Comparing the low-sulfide strain (*with* cysteine) and *E. coli* + Na_2_S (*without* cysteine), sulfide levels were similar (**Fig. 5c right**, *p* = 0.96), but thiosulfate levels were drastically different. This led us to hypothesize that the unexpectedly low thiosulfate concentration in the chip with the high-sulfide strain may instead be caused directly by the cysteine, either through an impact on bacterial/Caco-2 sulfur metabolism, or through abiotic reactions. For example, at high mM concentrations (~66 mM), cysteine and thiosulfate react slowly in aqueous conditions to produce cysteine and cysteine sulfonic acid^59^. If this reaction happens on-chip, this would explain the lower thiosulfate levels in the presence of cysteine. Resolving these intriguing possibilities is beyond the scope of this study. Nevertheless, use of the GMPS identified significant metabolic differences between the use of exogenous and microbially-produced sulfide. This is important because sulfide donors are the field standard^18–20,23,24,60,61^, yet cysteine-derived sulfide is likely the major source in the human gut^14^. Together, these results highlight the utility of the high-resolution view of metabolic crosstalk between the host and microbiota provided by our GMPS.

We next asked whether microbial H_2_S production could impact host gene expression in the GMPS. H_2_S is a known genotoxic agent and has been shown to change the transcriptional activity of various Caco-2 genes like growth arrest and DNA damage-inducible alpha (*GADD45a*)^19,43^. However, those experiments were conducted in static *in vitro* cultures with exogenous H_2_S sources. We therefore determined the Caco-2 *GADD45a* transcriptional response on-chip to different levels of microbially produced H_2_S by RT-qPCR (**Fig. 5d**). Caco-2 cells co-cultured with the high-sulfide strain for two days had a 6.2-fold increase in *GADD45a* expression compared to control (no sulfide). The high-sulfide strain produced 3.5-fold more H_2_S and caused a two-fold increase in Caco-2 *GADD45a* expression compared to the low-sulfide strain (**Fig. 5d**, *p* = 0.07). This effect is likely mediated by H_2_S-induced DNA damage leading to cell cycle arrest^19,43^, and supports the hypothesis that H_2_S affects the host in a concentration-dependent manner^11,20^. Importantly, the effect of exogenous Na_2_S on *GADD45a* expression is more variable than the high and low-sulfide strains, with coefficients of variation of 0.45, 0.32, and 0.14, respectively, highlighting the increased precision in dosing achievable by an engineered microbial source of H_2_S. Our system re-creates known H_2_S biological effects on Caco-2 cells^19,43^, further validating that this platform can be used to investigate how microbially derived metabolites affect human host cells.

## Discussion

Investigating the mechanistic role of microbially derived metabolites on host biology remains experimentally difficult. Specifically, the inability to accurately model human intestinal biology in 2D static cultures, coupled with the limited tools to control and measure gut metabolite levels *in vivo* make it difficult to accurately titrate relevant metabolites in a physiologically relevant experimental system. We posited that metabolically engineering microbes in a GMPS to produce the desired metabolite from resources available in the gut could represent a generalizable approach to address these challenges. In this proof-of-concept study, we focused on H_2_S owing to i) its debated role in human gut diseases, ii) the wide and uncertain range of physiologically relevant concentrations reported^10–12,14–18,20,51,54–56^, and iii) its volatility, which makes it challenging to study with conventional static cultures, where exogenous sources of H_2_S either evaporate quickly or are slowly released through chemical donors with unpredictable breakdown kinetics.

We used a synthetic biology approach to develop a small *E. coli* strain library that successfully titrated H_2_S across this wide range. Interestingly, we discovered a synergistic effect on H_2_S production when co-expressing the *E. coli* cysteine transporter (*yhaO*) and desulfidase (*yhaM)*, with this strain resulting in 210% more H_2_S than the total production from strains expressing these genes individually. This suggests that high H_2_S production is limited by the cell’s capacity both for cysteine transport and catalysis. Compared to the transport of other amino acids, uptake of cysteine is poorly characterized, and dedicated transporters are not widely distributed^63^. Because of this, it has been posited that the preferred source of sulfur for microbes in anoxic conditions like the gut is H_2_S, since it can freely diffuse across the cell membrane, allowing cells to avoid the energy investment in transport. As such, overexpression of the *yhaO* transporter in our strain may allow it to compete effectively for cysteine in a more complex microbiome, enabling its deployment in *in vivo* models.

Once in co-culture on the chip, the strains produced H_2_S across the gut physiological range. The low-sulfide strain, expressing a heterologous desulfidase, controllably produced H_2_S across a ~200 μM range through IPTG titration. This result showcases the high degree of control over microbially-derived H_2_S levels in our GMPS, which was unachievable with the exogenous sulfide donor, Na_2_S. Compared to microbially derived H_2_S, dosing exogenous Na_2_S resulted in a larger variance in H_2_S effluent measurements, effect on P_app_, and *GADD45a* mRNA expression. Na_2_S is not a natural sulfide source in the human gut, and HPLC analysis showed different thiosulfate profiles when compared to cysteine-derived sulfide, which is a major production pathway in the human gut^14^. Additionally, diffusion of H_2_S to the epithelial cell surface may depend on gut biogeography. For example, *Desulfibrio* spp., major H_2_S producers in the human gut, colonize and metabolize gut mucus^64^ and may produce local sulfide gradients which would not be replicated by small-molecule sulfide donors. Collectively, these results highlight the advantages of using engineered microbes over chemical donors for precise modulation of the gut metabolome.

There are several advantages to use of our GMPS platform. First, the system was designed with gas impermeable materials that retain gas tension better than PDMS^41,42^, specifically to study volatile metabolites. The experiments reported here would have been infeasible in PDMS-based GoC systems due to H_2_S evaporation. Second, compared to static co-culture with organoids, the constant fluid flow in the system promotes the stable co-culture of microbes and host epithelial cells, enabling longer duration experiments without the problems of microbial overgrowth^29,30^. We successfully maintained the system for 48 hours. There were no experimental factors that prevented co-cultures beyond two days (e.g. overgrowth, Caco-2 cell death), so future work could explore chronic sulfide effects on the host. Third, we engineered the system to be optically transparent, enabling real-time visualization of *E. coli* colonization without the need to terminate the experiment. The stable colonization of this chip is likely attributable to mucus production, as shown previously in our GMPS^41^. Finally, compared to mouse models, our GMPS permits higher-throughput experimentation, and its use of human tissue overcomes some of the limitations arising from different species^27^. Direct access to the lumen and apical material in the simplified system facilitates the experimental study of host-microbiota metabolic cross-talk, which is experimentally challenging in mice^39^.

One limitation in our study is the lack of a complex microbiota, which that may modulate the effects of sulfide on the host, and impact H_2_S production through competition for cysteine. A previous study using mice showed that introduction of a H_2_S-producing microbe, *Atopobium parvulum,* induced colitis only in the presence of an intact microbiota^12^. Thus, it would be interesting to examine the impact of controlled sulfide production on-chip using our strains in the presence of additional gut bacteria. This could be done in our system by taking advantage of the gas impermeability to establish oxygen gradients supportive of aerobic and anaerobic bacteria, as has been done in other GoC systems^35^. Regarding cysteine competition, as described above, the expression of a dedicated cysteine transporter may enhance the competitiveness of our strain. In addition, we could explore other pathways for H_2_S production. For example, *E. coli* can accumulate the tripeptide glutathione at very high levels intracellularly^65^. This could be mobilized into cysteine on-chip by controlled induction of specific peptidases, then converted to H_2_S as described here, providing a source of microbial sulfide inaccessible to other microbes. Additionally, as gut-related disorders involve an immune component, expanding the types of human tissue used in our GMPS could allow further interrogation of important aspects of the host response. While not inclusive of the biodiversity in the human gut, we note that the immortalized Caco-2 cell line used here was chosen to minimize the biological variability present in primary cultures, as the focus was on assessing the potential of engineered bacteria to manipulate metabolite levels. Future studies will incorporate additional cell types in order to further explore the biological implications of H_2_S production, and identify factors underlying the robustness of our engineered bacteria.

We believe the combination of engineered microbes and the GMPS is a powerful platform for addressing a range of important topics. The gas-impermeable nature of our platform could be leveraged to ask fundamental questions about the other microbiota-derived gasotransmitters, NO and CO. These gaseous mediators likely affect the stromal gut enteric nervous and immune systems^66–68^. Studying these gases in animal models is more complicated than soluble metabolites due to their volatility and rapid diffusion through tissue^69,70^. In our study, basal H_2_S was detected on the chip at concentrations similar to human serum^52^. This lays the foundational groundwork to incorporate vascular, neuronal, or immune cells in the basal compartment^32,34^ to investigate how microbiota-derived gasotransmitters affect the local tissue environment. This platform is not limited to gaseous metabolites. Synthetic biology allows for plug-and-play of different genetic elements to study other metabolites, such as neurotransmitters. Gut microbes affect the brain’s neurotransmitter pool by modulating precursors in the gut that communicate via the gut-brain-axis^71^.

More broadly, translating synthetic probiotics from test tubes to the clinic will require extensive validation and additional engineering to ensure robust performance in the complex gut environment. Incorporating the GMPS into the design-build-test-learn cycle for smart probiotics could help identify factors underlying strain stability, performance, and robustness at lower cost and with more fidelity than mouse studies. Critically, GoC systems have accurately predicted the *in vivo* strain activity of a clinical biotherapeutic^38^, supporting their role as an orthogonal pre-clinical model. Overall, we have validated a powerful platform combining synthetic biology and gas tension-retaining GMPSs for investigating the role of gaseous mediators in the human intestine. This platform enables precise manipulation of the gut metabolome, which we leveraged to understand the concentration-dependent effects of H_2_S on human gut epithelial cells. This platform will allow researchers to experimentally address host-microbe hypotheses that are infeasible with current animal and *in vitro* models.

## Methods

### Strain development and growth curves

The *decR*, *sseA*, and *malY* genes were knocked out of *E. coli* MG1655 using the λ-Red recombination and CRISPR Cas9 systems^49^. The pCas (Addgene plasmid #62225) and pTargetF (Addgene plasmid #62226) plasmids were gifts from Sheng Yang^49^. A custom N20 sequence was added to pTargetF via PCR to target each gene of interest directly. pCas and pTargetF were transformed and cured according to the protocol^49^. Colonies were picked, and knockouts were verified with PCR and Sanger sequencing (Genewiz from Azenta Life Sciences).

The strains were grown overnight in M9 minimal media (4 g/L glucose, 1 g/L casamino acids (cat. no 2240, VWR), and 1 mg/L thiamine (cat. no T4625, Sigma-Aldrich)) and diluted to an OD_600_ of 0.01 with M9 minimal media or M9 minimal media with 5 mM cysteine (cat no. 30129, Millipore Sigma). The strains were added to 96 well plates (cover on) and were grown for 16 hours at 37C in a plate reader (SpectraMax i3, Molecular Devices) in technical triplicates or quadruplicates. The plate was shaken at medium speed and OD_600_ reads were taken every ten minutes.

*E. coli* genes (*decR*, *yhaM*, *yhaO*) were cloned from *E. coli* MG1655 gDNA (cat no. T3010, New England Biolabs Inc.) via PCR. The *cdl* gene (from *F. nucleatum*) was codon optimized for *E. coli* and synthesized (Integrated DNA Technologies). Gibson assembly (cat no. E2621, New England Biolabs Inc.) was used to assemble backbones and genes of interest. Once assembled, plasmids were chemically transformed into DH5-alpha cells (cat no. C2987, New England Biolabs Inc.) and plated on agar plates containing appropriate antibiotics. Colonies were picked and were sequence verified with PCR and Sanger sequencing. Once verified, plasmids were mini-prepped (cat no. D4211, Zymo Research) and transformed into the chemically competent MG1655-Δ*decR*. Strain names and descriptions can be found in **Table S2**. All primers used in cloning and RT-qPCR can be found in **Table S3**.

### Hungate tube experiments

Cultures were grown overnight in M9 minimal media and diluted 1:25 with M9 minimal media in shake flasks. For strains bearing plasmids with the lacUV5 (variable induction), T7 (100 μM IPTG induction), and pBAD (10 mM L-arabinose induction) promoters, they were induced at OD_600_ 0.6 for two hours. Cultures were centrifuged and resuspended to OD_600_ 0.4 with PBS++ (non-growth experiments) or M9 minimal media (growth experiments). Negative controls had no cysteine. 5 mL of culture were added to sterile Hungate tubes and placed in a shaking incubator at 37C. Samples were taken with sterile needles at hours two and four while maintaining a closed and sterile system. H_2_S production was quantified with the methylene blue assay.

### H_2_S detection by methylene blue assay

The protocol was adapted and adjusted to detect H_2_S in liquid samples^72^. 200 μL of the experimental sample was added to 615 μL zinc acetate mixture (600 μL of 1% w/v zinc acetate dihydrate with 15 uL 3 M NaOH) and vortexed. After 5-10 minutes of incubating at room temperature, 150 μL of 0.1% n-n-dimethylethylenediamine (cat no. D157805, Millipore Sigma) in 5 M hydrochloric acid was added, followed by 150 μL of 23 mM ferric chloride (cat no. 12321, Sigma Aldrich) in 1 M hydrochloric acid. Samples were centrifuged at 16,000 RCF for 2 minutes. 200 μL of supernatant were plated in a transparent 96-well plate and were quantified using absorbance (670 nm) (SpectraMax i3, Molecular Devices).

For Hungate tube experiments, 200 μL samples were taken from Hungate tubes with a sterile syringe needle while maintaining a sterile closed system. For GMPS experiments, apical and basal effluent tubing was fed into collection tubes containing 2.2 mL of zinc acetate mixture to trap H_2_S over the sampling period (four hours at 180 μL/min). After four hours, effluent collection tubes were vortexed, and 815 μL of effluent/zinc acetate mixture was taken to be used for subsequent steps.

Standard curves were made by dissolving Na_2_S in 0.1 M NaOH and serially diluted in PBS++ or DMEM. The zinc acetate mixture was immediately added, tubes were vortexed, and samples sat at room temperature for at least 5 minutes. Mixtures were used for subsequent steps, and methylene blue was measured as described. Sample standard curves of PBS++ and DMEM are provided in **Supp. Fig. 5**.

### GMPS fabrication

The GMPS device used in this study is a derivative of previous work from our lab^41^. The fabrication used poly(methyl methacrylate) (PMMA) sheets, PET membranes, double-sided adhesive tape, and a laser cutter. Each layer of the chip was designed with computer-aided drafting (CAD, Autodesk Inventor) and was laser cut (Epilog Zing 16, Epilog Laser). The 3/16” PMMA (PMMA, McMaster-Carr) contained six laser cut holes, two for inlet flow, two for outlet flow, and two for access to apical and basal channels (referred to as ‘seed ports’), which were tapped to create threading for Luer lock fittings. This piece served as the top layer of the device. The second layer was a sheet of 1/16” PMMA (PMMA, McMaster-Carr) inserted between two pieces of 50 μm thick double-sided adhesive tape (966 Adhesive Transfer Tape, 3M), which served as the ceiling of the apical channel. All six circular inlets and outlets were laser-cut to match the through-holes in the top layer. The third layer was a PET membrane with 0.4 μm diameter pores (ipCELLCULTURE Track Etched Membrane, it4ip S.A.) that sat between the second and fourth layers, creating a cell culture surface for the apical channel. Circular inlets and outlets were laser-cut to match the through-holes in the fourth layer. A sheet of 1/16” PMMA inserted between two pieces of 50 μm thick double-sided adhesive tape served as the fourth layer and the ceiling of the basal channel. Three circular inlets and outlets were laser cut to match the through-holes in the top two layers. The fifth layer was a no. 1 glass coverslip that sealed the basal channel. Using a custom device, the layers were pressed together while maintaining the alignment of the holes and channels.

After assembly, the chips were pressed and stored under vacuum at 50C for two days to rid gas bubbles in the device. Male Leur lock fittings (cat no 40064, QOSINA) with Viton O-Rings 5/32 “ID 9/32” OD (cat no. 1284N107, McMaster-Carr) were fitted to all six threaded holes in the 3/16” layer. Female Leur locks to barb connectors were fitted to the two inlets and outlets to connect Soft PVC Clear Tubing, 1/16 “ID, 1/8” OD (McMaster-Carr). The apical port served as a seed port for Caco-2 and bacteria. Otherwise, throughout the experiment, the two seed ports were capped and sealed. The PVC tubing was connected to a syringe pump (NE-1600 Six Channel Programmable Syringe Pump, New Era Pump Systems Inc.), perfusing culture medium through the chip. Details of the GMPS can be found in **Table S1** and a schematic in **Figure 1**.

The fluidic shear stress (τ, dyne/cm^2^) was calculated by the approximation of flow between two parallel plates^36,42^

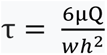

where μ is the dynamic viscosity of the apical fluid^40^ [0.0097 g/cm-s], Q is the volumetric flow rate [0.00005 cm^3^/sec], w is the width of the apical channel [0.1 cm], and h is the height of the apical channel [0.0196 cm].

### Caco-2 cell culture in GMPS

Caco-2 cells were obtained from the American Type Culture Collection (ATCC) and cultured in Dulbecco’s Modified Eagle Medium (DMEM, cat no. MT10013CV, Corning) supplemented with 10% fetal bovine serum (FBS, cat no. 10437028, ThermoFisher) and 100 U/mL penicillin−streptomycin (cat no. 15140122, ThermoFisher).

Microfluidic chips were removed from the vacuum chamber and UV sterilized (300 mJ/cm^2^) for 10 minutes (Spectrolinker XL-1000, Spectronics Corporation) on each side. Chips were then treated with oxygen plasma (Expanded Plasma Cleaner PDC-001, Harrick Plasma) for 1 minute. Fittings and tubing were sterilized by autoclaving. The chips, fittings, tubing, and syringe pump were assembled in a biosafety cabinet using an aseptic technique. The apical channel was coated in a 400 μg/mL solution of rat tail type I collagen (cat no. 354249, Corning), after which the chips were placed in a humidified incubator at 37C and 5% CO_2_ for one hour. After this time, the culture medium was perfused through the chip in preparation for seeding Caco-2. Caco-2 cells were harvested from flasks and concentrated to 5,000,000 cells/mL. 200 μL of Caco-2 cells were manually injected into the chip via the apical seed port. The chip set-up was placed in a humidified incubator at 37C and 5% CO_2_ without flow for two hours to allow cell adhesion. After two hours, the apical and basal flow was initiated at 3 μL/minute (shear stress 0.076 dyne/cm^2^). Effluent media was collected in waste collection tubes. Chips were under flow for seven days. On the seventh day, apical media was swapped for sterile PBS++ (pH 7.4) with 25 mM Lucifer Yellow (cat no. L453, Thermo Fisher). PBS++ was made of 1 L of DPBS, no calcium, no magnesium (cat no. 14190144, Thermo Fisher), 2.4 g/L HEPES (cat no. BP310-1, Fisher Scientific), 100 g/L anhydrous calcium chloride (cat no. 349615000, Thermo Fisher), and 48.4 g/L pure magnesium chloride (cat no. 223210010, Thermo Fisher). The basal media was swapped with antibiotic-free DMEM with 10% FBS in preparation for the inoculation of engineered strains. The media was allowed to perfuse through the chip overnight to clear residual antibiotics. For this study, Caco-2 cells were between passage numbers 25 and 47.

### Inoculating bacteria in GMPS

One to two hours before inoculation, the GMPS set-up was moved to the biosafety cabinet, and all apical and basal syringes were swapped to prime the chip. Before priming the chip, the media was filter sterilized. The basal media was swapped with antibiotic-free DMEM with 10% FBS in preparation for the inoculation of engineered strains. The following table describes the contents of each apical syringe used for each experiment. The syringes were flowed at 30 μL/min for ten minutes to clear any air bubbles introduced during handling. After ten minutes, the flow was restored to 3 μL/min.

| Experimental name | Syringe content description |
| --- | --- |
| Control | PBS++, 25 mM Lucifer Yellow, 5 mM L-cysteine, 250 μM IPTG, 50 μg/mL chloramphenicol. Media pH 7.4. |
| DSS | PBS++, 25 mM Lucifer Yellow, and 3% w/v DSS. Media pH 7.4. |
| Na<sub>2</sub>S | PBS++, 25 mM Lucifer Yellow, and 5 mM Na<sub>2</sub>S. Media pH 7.4. |
| Lac-*cdl* | PBS++, 25 mM Lucifer Yellow, 5 mM L-cysteine, variable IPTG (0, 2.5, or 250 μM), and 50 μg/mL chloramphenicol. Media pH 7.4. |
| T7-*yhaM* + BAD-*yhaO* | PBS++, 25 mM Lucifer Yellow, 5 mM L-cysteine, 100 μM IPTG, 10 mM L-arabinose, 50 μg/mL chloramphenicol, and 50 μg/mL spectinomycin. Media pH 7.4. |
| T7-*yhaM* + BAD-*yhaO* (no cysteine) | PBS++, 25 mM Lucifer Yellow, 100 μM IPTG, 10 mM L-arabinose, 50 μg/mL chloramphenicol, and 50 μg/mL spectinomycin. Media pH 7.4. |
| T7-*yhaM* + BAD-*yhaO* (no cysteine) + Na<sub>2</sub>S | PBS++, 25 mM Lucifer Yellow, 5 mM Na<sub>2</sub>S, 100 μM IPTG, 10 mM L-arabinose, 50 μg/mL chloramphenicol, and 50 μg/mL spectinomycin. Media pH 7.4. |

Bacterial cultures were grown overnight in M9 minimal media and diluted 1:25 with M9 minimal media in shake flasks. For strains bearing plasmids with the lacUV5 (variable induction 0-250 μM IPTG), T7 (100 μM IPTG induction), and pBAD (10 mM L-arabinose induction) promoters, they were induced at OD_600_ 0.6 for two hours. Cultures were centrifuged and resuspended to OD_600_ 0.4 with their respective apical channel media. Sodium sulfide nonahydrate (Na_2_S) (cat no. 208043, Sigma Aldrich) was used for exogenous sulfide. DSS (cat no. AC433240050 Fisher Scientific) was dissolved in PBS++ to 3% w/v.

Next, the GMPS set-up was moved to the biosafety cabinet, and flow was paused. 200 μL of bacteria were injected into the apical seed port at an OD_600_ of 0.4 using a syringe pump (100 μL/min). Once inoculated, the chip set-up was moved back into the humidified incubator at 37C 5% CO_2_, and flow was resumed immediately at 3 μL/min. Throughout this study, inoculation is defined as hour zero.

### Gut permeability analysis (P_app_ calculations)

25 mM Lucifer Yellow was added to apical syringes the night before inoculating the chips. Four hours prior to inoculation, GMPS basal effluent was collected for two hours to establish a baseline P_app_. Additionally, effluent samples were collected at hours 18 and 42 of the experiment. Samples were quantified using a DMEM standard curve using a fluorescence plate reader (SpectraMax i3, Molecular Devices) (ex: 485 nm, em: 535 nm). The CO_2_ in the incubator drives down the pH of DMEM, affecting the fluorescent properties at these excitation/emission values. To control for CO_2_-induced pH changes in the DMEM, an additional DMEM syringe was incubated overnight and used to make the standard curve. Apparent permeability (P_app_, cm/sec) was calculated with the following equation

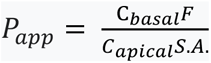

where C_basal_ is the concentration of Lucifer Yellow detected in the basal channel as determined by the standard curve [μM], F is the flow rate [0.00005 cm^3^/sec], C_apical_ is the concentration of Lucifer Yellow in the apical channel [25 μM], and S.A. is the surface area of the Caco-2 monolayer [0.1341 cm^2^].

### CFU analysis

Apical and basal effluent was collected at hours −4, 18, and 42. Apical samples were serially diluted 1,000-fold or 10,000-fold, and 50 μL were plated on LB agar plates containing appropriate antibiotics or antibiotic-free plates for control chip samples. For basal samples, 50 μL of basal effluent was directly plated. Agar plates were incubated overnight at 37C and counted the following day. For apical samples, the CFU count was multiplied by a factor of 20,000 (for 1,000-fold diluted samples) or 200,000 (for 10,000-fold diluted samples) to get units of CFU/mL. In bacteria-free chips, apical and basal effluent were plated on LB agar plates, and no colonies were detected.

### Metabolite quantification with HPLC

Apical and basal effluent samples were collected at hours −4, 18, and 42 and was immediately stored at −80C. For glucose and acetate analysis, samples were thawed, centrifuged, and the supernatant was loaded into HPLC tubes. An Agilent 1260 HPLC Infinity II system was used with an isocratic flow of 14 mM sulfuric acid at 0.6 mL/min. 10 μL of the sample were injected into an Aminex HPX 87H ion (300×7.8 mm) C18 reversed phase column (cat no. 1250140, BioRad) at 60C fitted with a Micro-Guard Cation H cartridge (cat no. 1250129, Bio-Rad). Compounds were identified with a refractive index detector (1260 Infinity II Refractive Index Detector, Agilent) at 50C. Calibration curves of the pure compounds were used to identify and quantify compounds in experimental samples. The protocol was adapted and adjusted from the previous work^73^.

For detecting thiosulfate, samples were thawed, centrifuged, and the supernatant was used for monobromobimane (mBBr) derivatization (cat no. 69898, Millipore Sigma). To derivatize samples, 5 mM mBBr was dissolved in 200 mM Tris-HCl. 10 μL of supernatant were added to 90 μL mBBr solution and incubated at room temperature for one hour, and samples were covered in foil. After one hour, 30 μL of 1M hydrochloric acid was added to terminate the reaction. Samples were centrifuged, and the supernatant was placed in HPLC tubes. 5 μL of the sample were injected into a C18 reversed phase column (InfinityLab Poroshell 120 EC-C18 2.7 μm 3×100 mm, Agilent) at 40C and detected with an HPLC fluorescence detector (1260 Infinity II Fluorescence Detector, Agilent) at an excitation/emission spectra of 384/478 nm. The flow rate through the column was 0.5 mL/min, and a gradient elution was applied to separate thiols: 0-2 min, 0% B; 2-8 min, 46% B; 8-9 min, 64% B; 9-13 min, 100% B; 13-14 min, 0% B; 14-15 min, 0% B. Solvent A was 10% methanol and 0.25% glacial acetic acid, adjusted to pH 3.9. Solvent B was 90% methanol and 0.25% glacial acetic acid, adjusted to pH 3.9. The protocol was adapted and modified from previously described protocols^74,75^.

### RT-qPCR analysis

After two days of co-culture, the flow was stopped, and the GMPS set-up was removed from the incubator. Via the seed port, PBS++ was used to wash the apical channel, and 300 μL RNA lysis buffer (cat no. T2010, New England Biolabs Inc.) was applied to the channel and vigorously pipetted. RNA was purified following the manufacturer’s protocol (cat no. T2010, New England Biolabs Inc.). Oligo(dT)_12-18_ (cat no. 18418012, Thermo Fisher) was used to reverse transcribe Caco-2 RNA (C1000 Touch Thermal Cycler, BioRad). The resulting cDNA was cleaned and concentrated (cat no. D40144, Zymo Research) and 1 μL was used for qPCR. A SYBR Green-based qPCR master mix (cat no. M3003, New England Biolabs Inc.) and primers (**Table S3**.) were used to amplify genes of interest (C1000 Thermal Cycler with CFX96 Real-Time System, BioRad). Primers were made with NCBI Primer Blast^76^. The target gene, *GADD45a*, was normalized to *GAPDH*. Relative expression differences were calculated using the delta-delta Ct method.

### Immunofluorescent imaging in GMPS

Images of tight junction formation were assessed using fluorescence microscopy. Monolayers were fixed by flowing 4% formaldehyde (v/v in 2.5% Goat Serum) at 9 μL/min for 30 minutes, 0.1% TritonX (v/v in 2.5% Goat Serum) at 9 μL/min for 20 minutes, and 2.5% goat serum for 30 minutes at room temperature. Chips were covered in foil and placed in the fridge (4C) overnight. The basal channel was flushed with PBS. Flow rates were kept below 10 μL/min in the apical channel to prevent high shear stress and monolayer disruption. The following day, 1:200 ZO-1 conjugated antibody (cat no. MA3-39100-A647, Thermo Fisher) and 1:10,000 Hoechst (cat no. 62249, Thermo Fisher) were mixed in 2.5% goat serum and flowed at 6 μL/min for one hour. After one hour, the flow was paused, and the system sat static for one hour. Afterward, the monolayers were washed with DPBS for three hours at 4 μL/min. The GMPS was imaged with a confocal microscope (Zeiss LSM 880 NLO) using the DAPI and Alexa Fluor 647 filters.

For live staining, T7-*yhaM-RFP* + BAD-*yhaO* was inoculated on the chip. 1:2,000 Hoechst was added to apical and basal channels and flowed on the chip at 3 μL/min for three hours. Adding a higher concentration of Hoechst to apical and basal channels enhanced the signal, which may be attributed to the sizeable microbial population in the apical channel disrupting the mass transport of Hoechst to the Caco-2 nuclei. Before imaging, the basal channel was flushed with PBS. Next, the GMPS was imaged with confocal microscopy using the DAPI and mRFP-1.2 filters.

A live video of the co-culture was captured using an inverted fluorescence microscope (Zeiss Axio Observer Z1). To do this, the GMPS was kept in an incubated microscope set up (37C and 5% CO_2_), and normal experimental conditions were maintained including perfusion through the chip. Images were captured overnight and assembled in a movie using Zen software (Zeiss) and Adobe Premier Pro. Image processing was done in Zen software (Zeiss). Gaussian smoothing was used for all images.

## Statistical analysis

A one-way ANOVA or two-way ANOVA followed by post hoc Tukey-Kramer analysis were used to determine statistical significance (*p < 0.05). Bar graphs represent the average value from independent experiments, and dots represent the result from each independent experiment. Error bars represent the ± standard deviation. GraphPad Prism was used to carry out the analysis. Graphs were formatted in Adobe Illustrator. For each independent experiment, bacterial cultures were grown from cryostock, and Caco-2 cells were harvested from flasks. Independent experiments were performed on different days. For GMPS experiments, each independent experiment represents an individual GMPS. For samples that were lower than the limit of quantification, they were labeled as not detected (Source Data File).

## Data Availability

All data generated in this study are available in the Source Data file provided in this paper. The datasets generated during the current study are available from the corresponding authors upon reasonable request.

## Acknowledgments

We thank McKayla Van Orden for help cloning constructs, Ryan Brady for the CAD drawings of the microfluidic device, and Adam Bindas for help with the GMPS workflow development. Also, Robert Egan for allowing us to use the Northeastern University Department of Chemical Engineering machine shop.

In addition, we thank the Institute for Chemical Imaging of Living Systems at Northeastern University for consultation and imaging support and the Discovery cluster, supported by Northeastern University’s Research Computing team.

This material is based upon work supported by the National Science Foundation Graduate Research Fellowship under Grant Number 1938052 to J.A.H., and by the National Institute of Biomedical Imaging and Bioengineering of the National Institutes of Health under award number R21EB033892 to B.M.W. Any opinions, findings, conclusions, or recommendations expressed in this material are those of the authors and do not necessarily reflect the views of the funders.

## Authorship statement

J.A.H., A.N.K., R.K., and B.M.W. conceived the study and experimental design. J.A.H. conducted the experiments, collected and analyzed the data, and is the primary manuscript author. A.W.L. assisted in molecular cloning and data collection. A.S.S. helped build the GMPS and assisted in data collection. M.T.F. assisted in molecular cloning and protocol development. All authors reviewed and approved the final document.

## Correspondence

Inquiries and requests for more information should be addressed to R.K. and B.M.W.

## Supplemental information

See attached supplemental files.

## References

1. Shreiner A, Kao J, Y. V. The gut microbiome in health and in disease. Cell 31, 69–75 (2015).

2. Moser, G., Fournier, C. & Peter, J. Intestinal microbiome-gut-brain axis and irritable bowel syndrome. Wiener Medizinische Wochenschrift 168, 62–66 (2018).

3. Lavelle, A. & Sokol, H. Gut microbiota-derived metabolites as key actors in inflammatory bowel disease. Nat. Rev. Gastroenterol. Hepatol. 17, 223–237 (2020).

4. Kim, J. & Lee, H. K. Potential Role of the Gut Microbiome In Colorectal Cancer Progression. Front. Immunol. 12, 807648 (2022).

5. Pascal, V. et al. A microbial signature for Crohn’s disease. Gut 66, 813–822 (2017).

6. Jeffery, I. B. et al. An irritable bowel syndrome subtype defined by species-specific alterations in faecal microbiota. Gut 61, 997–1006 (2012).

7. Wang, T. et al. Structural segregation of gut microbiota between colorectal cancer patients and healthy volunteers. ISME J. 6, 320–329 (2012).

8. Koh, A. & Bäckhed, F. From Association to Causality: the Role of the Gut Microbiota and Its Functional Products on Host Metabolism. Mol. Cell 78, 584–596 (2020).

9. Gasaly, N., de Vos, P. & Hermoso, M. A. Impact of Bacterial Metabolites on Gut Barrier Function and Host Immunity: A Focus on Bacterial Metabolism and Its Relevance for Intestinal Inflammation. Front. Immunol. 12, 658354 (2021).

10. Metwaly, A. et al. Integrated microbiota and metabolite profiles link Crohn’s disease to sulfur metabolism. Nat. Commun. 11, 4322 (2020).

11. Singh, S. & Lin, H. Hydrogen Sulfide in Physiology and Diseases of the Digestive Tract. Microorganisms 3, 866–889 (2015).

12. Buret, A. G., Allain, T., Motta, J.-P. & Wallace, J. L. Effects of Hydrogen Sulfide on the Microbiome: From Toxicity to Therapy. Antioxid. Redox Signal. 36, 211–219 (2021).

13. Wallace, J. L., Motta, J. P. & Buret, A. G. Hydrogen sulfide: An agent of stability at the microbiome-mucosa interface. Am. J. Physiol. - Gastrointest. Liver Physiol. 314, G143– G149 (2018).

14. Braccia, D. J., Jiang, X., Pop, M. & Hall, A. B. The Capacity to Produce Hydrogen Sulfide (H2S) via Cysteine Degradation Is Ubiquitous in the Human Gut Microbiome. Front. Microbiol. 12, 705583 (2021).

15. Loubinoux, J., Bronowicki, J. P., Pereira, I. A. C., Mougenel, J. L. & Le Faou, A. E. Sulfate-reducing bacteria in human feces and their association with inflammatory bowel diseases. FEMS Microbiol. Ecol. 40, 107–112 (2002).

16. Gibson, G. R., Cummings, J. H. & Macfarlane, G. T. Growth and activities of sulphate-reducing bacteria in gut contents of healthy subjects and patients with ulcerative colitis. FEMS Microbiol. Lett. 86, 103–111 (1991).

17. Levine, J., Ellis, C. J., Furne, J. K., Springfield, J. & Levitt, M. D. Fecal hydrogen sulfide production in ulcerative colitis. Am. J. Gastroenterol. 93, 83–87 (1998).

18. Sakuma, S. et al. Hydrogen sulfide donor GYY4137 suppresses proliferation of human colorectal cancer Caco-2 cells by inducing both cell cycle arrest and cell death. Heliyon 5, e02244 (2019).

19. Xu, W., Watanabe, K., Mizukami, Y., Yamamoto, Y. & Suzuki, T. Hydrogen sulfide suppresses the proliferation of intestinal epithelial cells through cell cycle arrest. Arch. Biochem. Biophys. 712, 109044 (2021).

20. Hellmich, M. R., Coletta, C., Chao, C. & Szabo, C. The therapeutic potential of cystathionine β-Synthetase/hydrogen sulfide inhibition in cancer. Antioxidants Redox Signal. 22, 424–448 (2015).

21. Beaumont, M. et al. Detrimental effects for colonocytes of an increased exposure to luminal hydrogen sulfide: The adaptive response. Free Radic. Biol. Med. 93, 155–164 (2016).

22. Mottawea, W. et al. Altered intestinal microbiota-host mitochondria crosstalk in new onset Crohn’s disease. Nat. Commun. 7, (2016).

23. Wallace, J. L., Dicay, M., McKnight, W. & Martin, G. R. Hydrogen sulfide enhances ulcer healing in rats. FASEB J. 21, 4070–4076 (2007).

24. Wallace, J. L., Vong, L., McKnight, W., Dicay, M. & Martin, G. R. Endogenous and Exogenous Hydrogen Sulfide Promotes Resolution of Colitis in Rats. Gastroenterology 137, 569–578.e1 (2009).

25. Fan, Y. & Pedersen, O. Gut microbiota in human metabolic health and disease. Nat. Rev. Microbiol. 19, 55–71 (2021).

26. Chen, H. et al. A Forward Chemical Genetic Screen Reveals Gut Microbiota Metabolites That Modulate Host Physiology. Cell 177, 1217–1231 (2019).

27. Ingber, D. E. Is it Time for Reviewer 3 to Request Human Organ Chip Experiments Instead of Animal Validation Studies? Adv. Sci. 2002030, (2020).

28. Nguyen, T. L. A., Vieira-Silva, S., Liston, A. & Raes, J. How informative is the mouse for human gut microbiota research? DMM Dis. Model. Mech. 8, 1–16 (2015).

29. Zhang, J. et al. Coculture of primary human colon monolayer with human gut bacteria. Nat. Protoc. 16, 3874–3900 (2021).

30. Puschhof, J. et al. Intestinal organoid cocultures with microbes. Nat. Protoc. 16, 4633– 4649 (2021).

31. Kasendra, M. et al. Development of a primary human Small Intestine-on-a-Chip using biopsy-derived organoids. Sci. Rep. 8, 2871 (2018).

32. Beaurivage, C. et al. Development of a human primary gut-on-a-chip to model inflammatory processes. Sci. Rep. 10, 21475 (2020).

33. Kasendra, M. et al. Duodenum intestine-chip for preclinical drug assessment in a human relevant model. Elife 9, e50135 (2020).

34. Kim, H. J., Li, H., Collins, J. J. & Ingber, D. E. Contributions of microbiome and mechanical deformation to intestinal bacterial overgrowth and inflammation in a human gut-on-a-chip. Proc. Natl. Acad. Sci. U. S. A. 113, E7–E15 (2015).

35. Jalili-Firoozinezhad, S. et al. A complex human gut microbiome cultured in an anaerobic intestine-on-a-chip. *Nat*. Biomed. Eng. 3, 520–531 (2019).

36. Kim, H. J., Huh, D., Hamilton, G. & Ingber, D. E. Human gut-on-a-chip inhabited by microbial flora that experiences intestinal peristalsis-like motions and flow. Lab Chip 12, 2165–2174 (2012).

37. Nelson, M. T. et al. Evaluation of Human Performance Aiding Live Synthetically Engineered Bacteria in a Gut-on-a-Chip. ACS Biomater. Sci. Eng. https://doi.org/10.1021/acsbiomaterials.2c00774 (2022) doi:10.1021/acsbiomaterials.2c00774.

38. Nelson, M. T. et al. Characterization of an engineered live bacterial therapeutic for the treatment of phenylketonuria in a human gut-on-a-chip. Nat. Commun. 12, 2805 (2021).

39. Tovaglieri, A. et al. Species-specific enhancement of enterohemorrhagic E. coli pathogenesis mediated by microbiome metabolites. Microbiome 7, (2019).

40. Park, T. E. et al. Hypoxia-enhanced Blood-Brain Barrier Chip recapitulates human barrier function and shuttling of drugs and antibodies. Nat. Commun. 10, (2019).

41. Hosic, S. et al. Rapid Prototyping of Multilayer Microphysiological Systems. ACS Biomater. Sci. Eng. 7, 2949–2963 (2021).

42. Soucy, J. R. et al. Reconfigurable Microphysiological Systems for Modeling Innervation and Multitissue Interactions. Adv. Biosyst. 4, (2020).

43. Attene-Ramos, M. S. et al. DNA Damage and Toxicogenomic Analyses of Hydrogen Sulfide in Human Intestinal Epithelial FHs 74 Int Cells. Environ. Mol. Mutagen. 51, 304– 314 (2010).

44. Shimada, T., Tanaka, K. & Ishihama, A. Transcription factor DecR (YbaO) controls detoxification of L-cysteine in Escherichia coli. Microbiol. (United Kingdom) 162, 1698– 1707 (2016).

45. Wang, J. et al. Hydrogen Sulfide From Cysteine Desulfurase, Not 3-Mercaptopyruvate Sulfurtransferase, Contributes to Sustaining Cell Growth and Bioenergetics in E. coli Under Anaerobic Conditions. Front. Microbiol. 10, 2357 (2019).

46. Li, K. et al. Escherichia coli uses separate enzymes to produce H2S and reactive sulfane sulfur from L-cysteine. Front. Microbiol. 10, 298 (2019).

47. Mironov, A. et al. Mechanism of H2S-mediated protection against oxidative stress in Escherichia coli. Proc. Natl. Acad. Sci. U. S. A. 114, 6022–6027 (2017).

48. Luhachack, L., Rasouly, A., Shamovsky, I. & Nudler, E. Transcription factor YcjW controls the emergency H2S production in E. coli. Nat. Commun. 10, 2868 (2019).

49. Jiang, Y. et al. Multigene editing in the Escherichia coli genome via the CRISPR-Cas9 system. Appl. Environ. Microbiol. 81, 2506–2514 (2015).

50. Shatalin, K., Shatalina, E., Mironov, A. & Nudler, E. H2S: A Universal Defense Against Antibiotics in Bacteria. Science (80-.). 334, 986–990 (2011).

51. Padron, G. C. et al. Shear rate sensitizes bacterial pathogens to H2O2 stress. Proc. Natl. Acad. Sci. 120, (2023).

52. Bahadoran, Z. et al. Association between serum hydrogen sulfide concentrations and dysglycemia: a population-based study. BMC Endocr. Disord. 22, 79 (2022).

53. von Rosenvinge, E. C., O’May, G. A., Macfarlane, S., Macfarlane, G. T. & Shirtliff, M. E. Microbial biofilms and gastrointestinal diseases. Pathog. Dis. 67, 25–38 (2013).

54. Ijssennagger, N., van der Meer, R. & van Mil, S. W. C. Sulfide as a Mucus Barrier-Breaker in Inflammatory Bowel Disease? Trends Mol. Med. 22, 190–199 (2016).

55. Shin, W. & Kim, H. J. Intestinal barrier dysfunction orchestrates the onset of inflammatory host-microbiome cross-talk in a human gut inflammation-on-a-chip. Proc. Natl. Acad. Sci. U. S. A. 115, E10539–E10547 (2018).

56. Kumar, K. K. V, Karnati, S., Reddy, M. B. & Chandramouli, R. Caco-2 cell lines in drug discovery-an updated perspective. J. basic Clin. Pharm. 1, 63–69 (2010).

57. Jackson, M. R., Melideo, S. L. & Jorns, M. S. Human sulfide:Quinone oxidoreductase catalyzes the first step in hydrogen sulfide metabolism and produces a sulfane sulfur metabolite. Biochemistry 51, 6804–6815 (2012).

58. Cheng, H., Donahue, J. L., Battle, S. E., Ray, W. K. & Larson, T. J. Biochemical and Genetic Characterization of PspE and GlpE, Two Singledomain Sulfurtransferases of Escherichia coli. Open Microbiol. J. 2, 18–28 (2008).

59. Berestova, T. V. et al. NMR study of thiosulfate-assisted oxidation of L-cysteine. Mendeleev Commun. 33, 99–102 (2023).

60. Chen, S. W. et al. Protective effect of hydrogen sulfide on TNF-α and IFN-γ-induced injury of intestinal epithelial barrier function in Caco-2 monolayers. Inflamm. Res. 64, 789–797 (2015).

61. Zhao, H., Yan, R., Zhou, X., Ji, F. & Zhang, B. Hydrogen sulfide improves colonic barrier integrity in DSS-induced inflammation in Caco-2 cells and mice. Int. Immunopharmacol. 39, 121–127 (2016).

62. Blackler, R., Syer, S., Bolla, M., Ongini, E. & Wallace, J. L. Gastrointestinal-sparing effects of novel nsaids in rats with compromised mucosal defence. PLoS One 7, e35196 (2012).

63. Zhou, Y. & Imlay, A. Escherichia coli K-12 Lacks a High-Affinity Assimilatory Cysteine Importer. MBio 11, (2020).

64. Earley, H. et al. A preliminary study examining the binding capacity of Akkermansia muciniphila and Desulfovibrio spp., to colonic mucin in health and ulcerative colitis. PLoS One 10, e0135280 (2015).

65. Zhang, J. et al. Systematic manipulation of glutathione metabolism in Escherichia coli for improved glutathione production. Microb. Cell Fact. 15, (2016).

66. Jimenez, M. Hydrogen sulfide as a signaling molecule in the enteric nervous system. Neurogastroenterol. Motil. 22, 1149–1153 (2010).

67. Byrne, J. D. et al. Delivery of therapeutic carbon monoxide by gas-entrapping materials. 14, 1–9 (2022).

68. Fagone, P., Mazzon, E., Bramanti, P., Bendtzen, K. & Nicoletti, F. Gasotransmitters and the immune system: Mode of action and novel therapeutic targets. Eur. J. Pharmacol. 834, 92–102 (2018).

69. Haselden, W. D., Kedarasetti, R. T. & Drew, P. J. Spatial and temporal patterns of nitric oxide diffusion and degradation drive emergent cerebrovascular dynamics. PLoS Computational Biology vol. 16 (2020).

70. Mao, Q. et al. Sensitive quantification of carbon monoxide in vivo reveals a protective role of circulating hemoglobin in CO intoxication. *Commun*. Biol. 4, 1–15 (2021).

71. Chen, Y., Xu, J. & Chen, Y. Regulation of neurotransmitters by the gut microbiota and effects on cognition in neurological disorders. Nutrients 13, 2099 (2021).

72. Li, Z.-G. Quantification of Hydrogen Sulfide Concentration Using Methylene Blue and 5,5 0-Dithiobis(2-Nitrobenzoic Acid) Methods in Plants. Methods Enzymol. 554, (2015).

73. Woolston, B. M., Emerson, D. F., Currie, D. H. & Stephanopoulos, G. Rediverting carbon flux in Clostridium ljungdahlii using CRISPR interference (CRISPRi). Metab. Eng. 48, 243–253 (2018).

74. Kurpet, K., Glowacki, R. & Grazyna, C. Simultaneous Determination of Human Serum Albumin and Low-Molecular-Weight Thiols after Derivatization with Monobromobimane. Molecules 26, 3321 (2021).

75. Luebke, J. L. et al. The CsoR-like sulfurtransferase repressor (CstR) is a persulfide sensor in Staphylococcus aureus. Mol. Microbiol. 94, 1343–1360 (2014).

76. Ye, J. et al. Primer-BLAST: A tool to design target-specific primers for polymerase chain reaction. BMC Bioinformatics 13, (2012).

