## Supplemental Information for "Engineered bacteria titrate hydrogen sulfide and induce concentration-dependent effects on host in a gut microphysiological system"

1. Department of Chemical Engineering
Northeastern University
360 Huntington Ave.
Boston, MA 02115, USA

2. Department of Mechanical and Industrial Engineering
Northeastern University
360 Huntington Ave.
Boston, MA 02115, USA

3. Department of Bioengineering
Northeastern University
360 Huntington Ave.
Boston, MA 02115, USA

**Supplementary Figures**

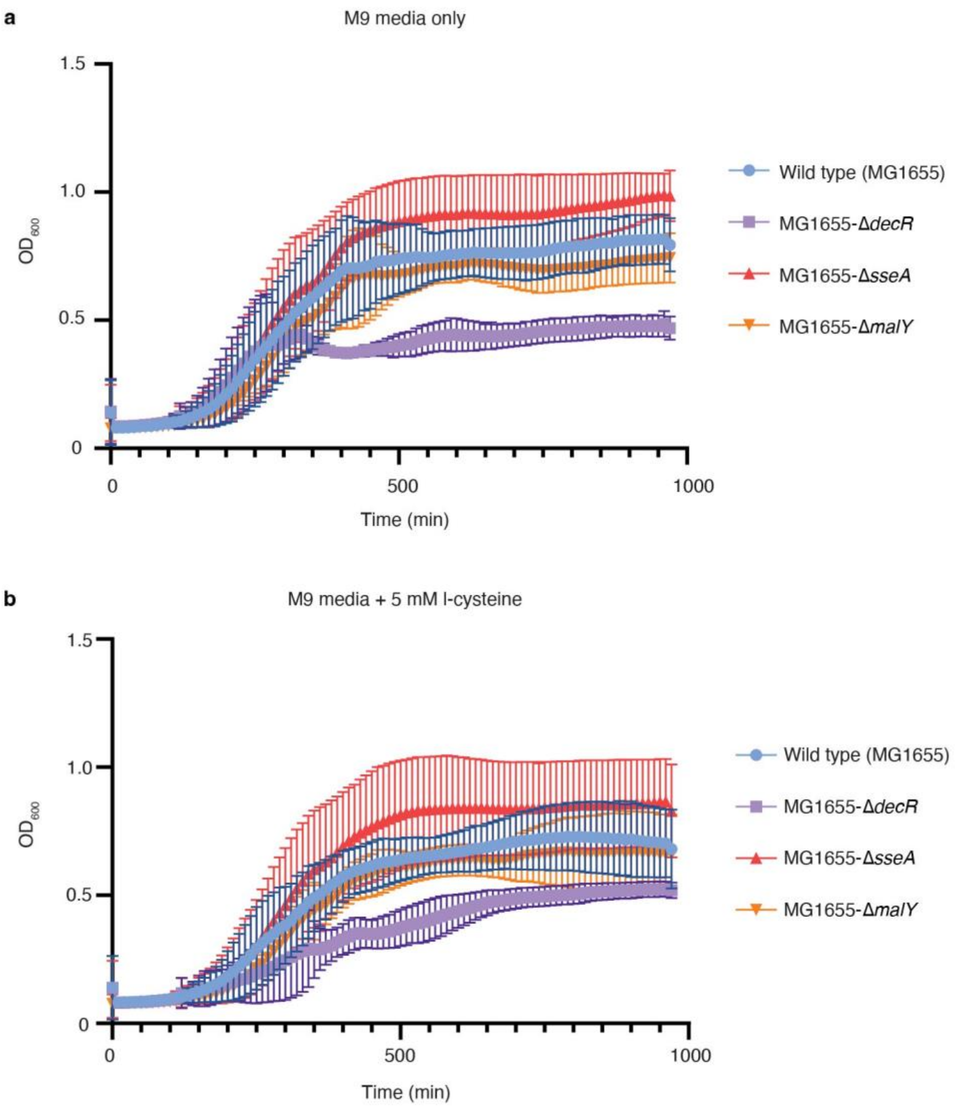


**Supplementary Figure 1. Characterization of gene knockouts on cell growth.**
**a-b** $OD_{600}$ readings were taken every 10 minutes for 16 hours. Each point is an average of 3 or 4 technical replicates. **a** $OD_{600}$ measurements of strains in M9 minimal media and **b** in M9 minimal media with 5 mM cysteine. n = 4 independent experiments, for *malY* n = 3 independent experiments. Source data can be found in the Source Data File.

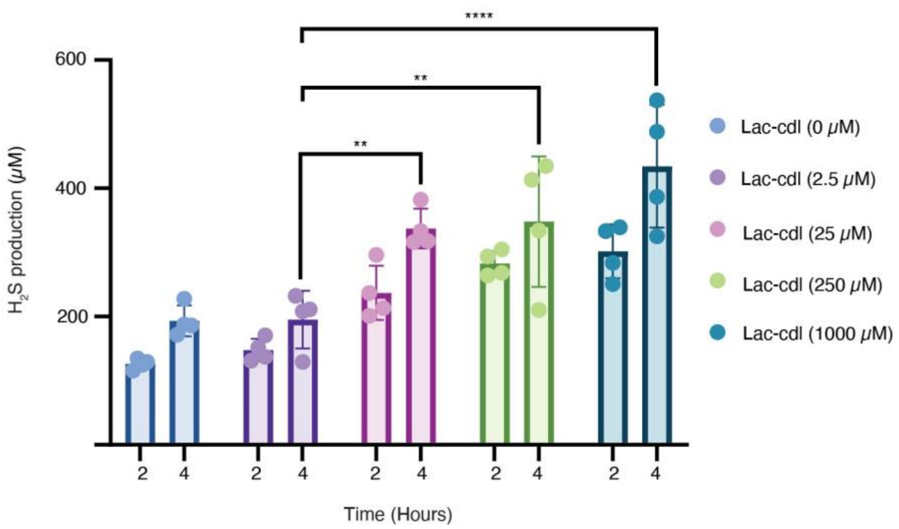


**Supplementary Figure 2. Lac-*cdl* titrates $H_2S$ in an IPTG-dependent manner in Hungate tubes.** Strains were grown and induced at different IPTG levels in shake flasks, spun down, diluted in PBS++ to $OD_{600}$ 0.4, and 5 mM cysteine was added. Samples were taken at hours two and four while maintaining a closed system. n = 4 independent experiments. Error bars represent SD and bars represent the mean value. **p < 0.01, ***p < 0.001, and ****p < 0.0001. 2-way ANOVA with post hoc Tukey analysis using a 95% confidence interval. Source data can be found in the Source Data File.

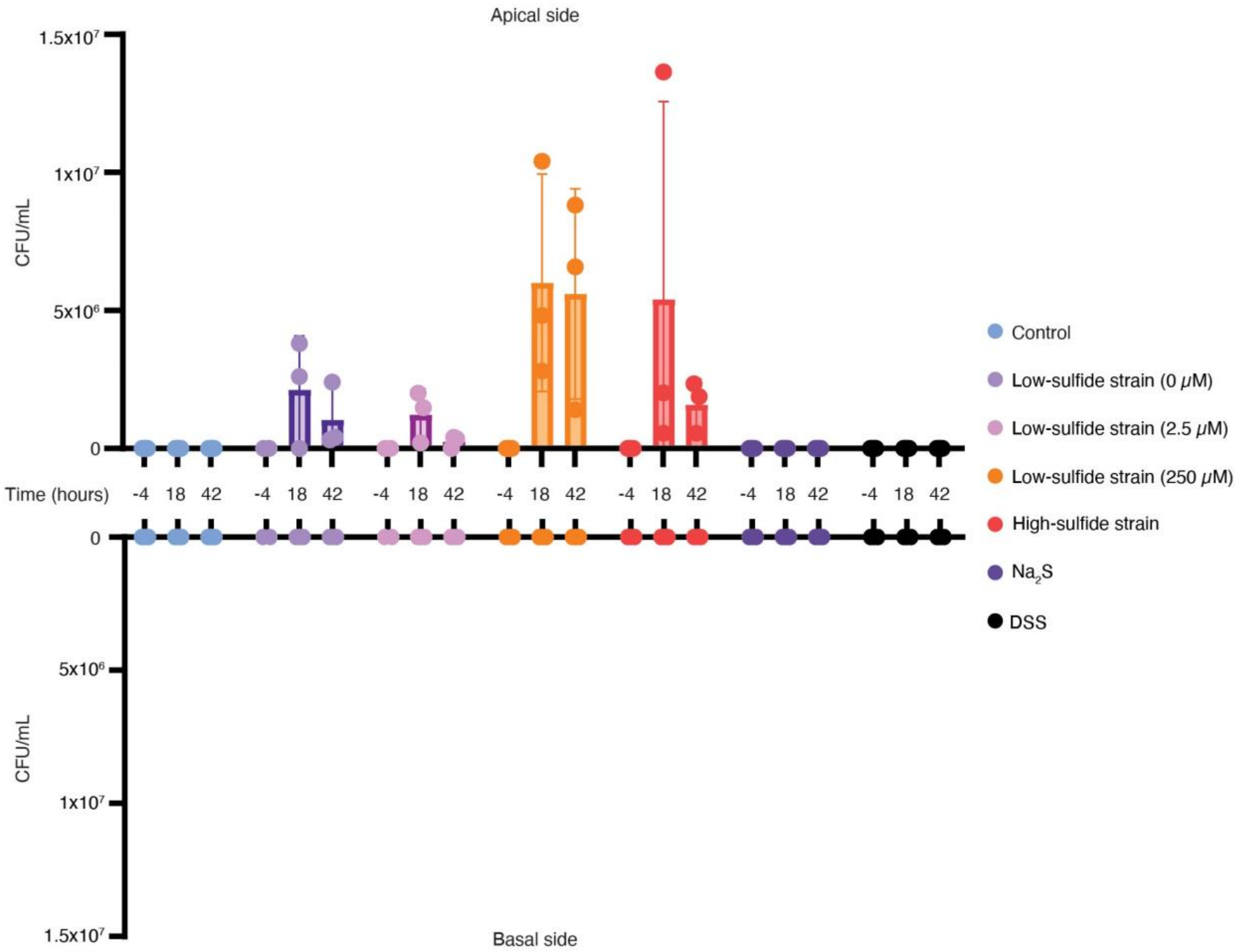


**Supplementary Figure 3. Quantification of microbial density in the GMPS.**
Apical and basal effluent media from the GMPS were plated and incubated overnight. CFUs were counted the following day to get a CFU/mL value. The top half of the graph represents the apical channel, and the bottom half represents the basal channel. For control, n = 4 independent experiments, for the other experimental conditions n = 3 independent experiments. Source data can be found in the Source Data File.

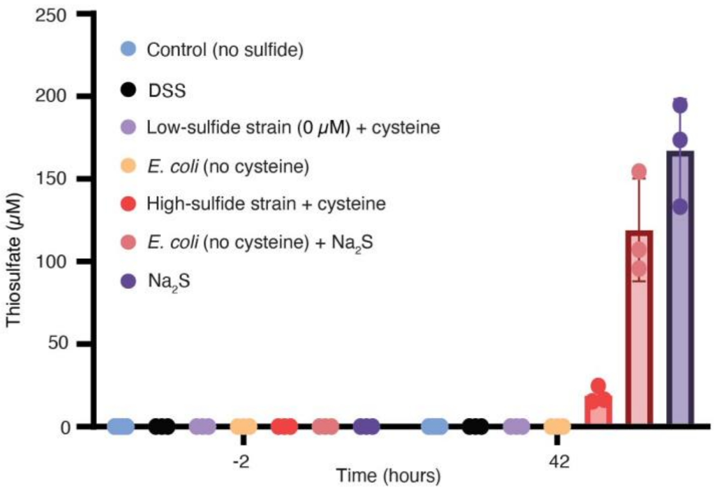


**Supplementary Figure 4. Thiosulfate detection from GMPS experiments.**
Apical effluent was collected and analyzed for thiosulfate via HPLC. n = 3 independent experiments, for Control; n = 4 independent experiments. Source data can be found in the Source Data File.

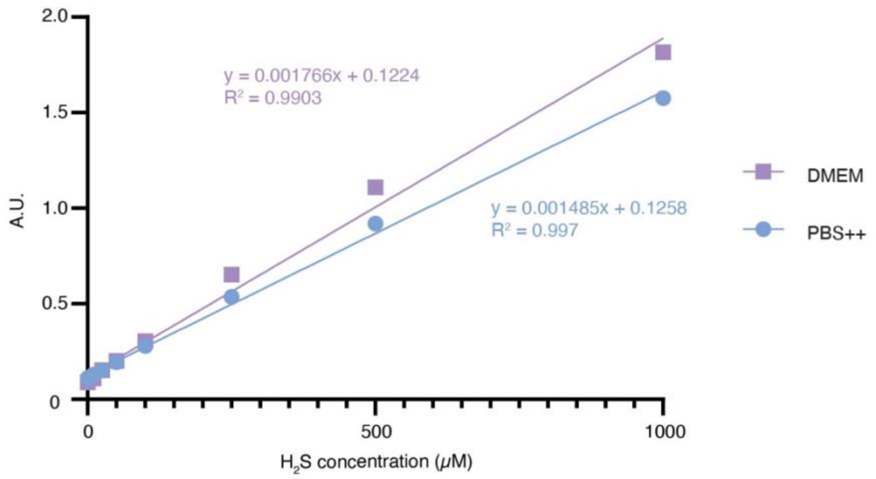


**Supplementary Figure 5. Example of $H_2S$ standard curves in PBS++ and DMEM.** Standard curves were made according to the methylene blue assay. Source data can be found in the Source Data File.

### Supplementary Tables

**Table S1. Chip features**

| **Chip feature** | **Value** |
|---|---|
| Apical flow | 3 µL/min |
| Basal flow | 3 µL/min |
| Shear stress | 0.076 dyne/cm$^2$ |
| Length of culturable channel | 1.341 cm |
| Width of culturable channel | 0.1 cm |
| Culturable surface area | 0.1341 cm$^2$ |
| Bacteria bolus concentration | $OD_{600}$ 0.4 |
| Bacteria bolus volume | 200 µL |
| Bacteria bolus injection speed | 100 µL/min |

**Table S2. Table of strains and their descriptions.**

| Strain name | Chassis strain | # of plasmids | Promoter-gene |
|---|---|---|---|
| Lac-*cdl* | MG1655-Δ*decR* | 1 | lacUV5-*cdl* |
| T7-*yhaM* | MG1655-Δ*decR* | 1 | T7-*yhaM* |
| BAD-*yhaO* | MG1655-Δ*decR* | 1 | pBAD-*yhaO* |
| T7-*yhaM* + BAD-*yhaO* | MG1655-Δ*decR* | 2 | T7-*yhaM* (plasmid 1)<br>pBAD-*yhaO* (plasmid 2) |
| T7-*yhaM-RFP* + BAD-*yhaO* | MG1655-Δ*decR* | 2 | T7-*yhaM* (plasmid 1)<br>cat-*RFP* (plasmid 1)<br>pBAD-*yhaO* (plasmid 2) |
| T7-*decR* | MG1655-Δ*decR* | 1 | T7-*decR* |
| T7-*cdl* | MG1655-Δ*decR* | 1 | T7-*cdl* |
| T7-*cdl* + BAD-*yhaO* | MG1655-Δ*decR* | 2 | T7-*cdl* (plasmid 1)<br>pBAD-*yhaO* (plasmid 2) |

**For Table S3, see the attached Excel file, “Source Data”.**

**For Supplemental Movie 1, see the attached file, “Supplemental Movie 1”.**
